# Expansion and accelerated evolution of 9-exon odorant receptors in *Polistes* paper wasps

**DOI:** 10.1101/2020.09.04.283903

**Authors:** Andrew W. Legan, Christopher M. Jernigan, Sara E. Miller, Matthieu F. Fuchs, Michael J. Sheehan

**Author notes:** Authors for Correspondence: Andrew W. Legan; Michael J. Sheehan.

## Abstract

Independent origins of sociality in bees and ants are associated with independent expansions of particular odorant receptor (OR) gene subfamilies. In ants, one clade within the OR gene family, the 9-exon subfamily, has dramatically expanded. These receptors detect cuticular hydrocarbons (CHCs), key social signaling molecules in insects. It is unclear to what extent 9-exon OR subfamily expansion is associated with the independent evolution of sociality across Hymenoptera, warranting studies of taxa with independently derived social behavior. Here we describe odorant receptor gene family evolution in the northern paper wasp, *Polistes fuscatus*, and compare it to four additional paper wasp species spanning ~40 million years of divergence. We find 200 functional OR genes in *P. fuscatus* matching predictions from neuroanatomy, and more than half of these are in the 9-exon subfamily. Lineage-specific expansions of 9-exon subfamily ORs are tandemly arrayed in *Polistes* genomes and exhibit a breakdown in microsynteny relative to tandem arrays in other OR subfamilies. There is evidence of episodic positive diversifying selection shaping ORs in expanded subfamilies, including 9-exon, E, H, and L, but 9-exon ORs do not stand out as selectively diversified among *Polistes* species. Accelerated evolution has resulted in lower amino acid similarity and high *d*N/*d*S among 9-exon ORs compared to other OR subfamilies. Patterns of OR evolution within *Polistes* are consistent with 9-exon OR function in CHC perception by combinatorial coding, with both selection and drift contributing to interspecies differences in copy number and sequence.

## INTRODUCTION

Odorant receptors are among the largest gene families in many animal genomes, with variation in the relative number of genes thought to reflect aspects of species chemosensory ecology. From the standpoint of molecular evolution, the odorant receptors (ORs) of insects and olfactory receptors in vertebrates have been widely studied as a model to understand the dynamics of gene family evolution (Nozawa & Nei 2007; Nei 2013; Benton 2015). Yet fundamental features of odorant receptor evolution remain unclear – why do some groups show predominantly conserved OR repertoires across species while others show rapid turnover in gene content or accelerated rates of evolution? Moreover, the relative importance of social interactions, sexual selection, and ecology in shaping patterns of OR evolution within and between clades is poorly understood. Comparative studies of distantly related species have provided insights into the evolutionary processes shaping the OR gene family at broad phylogenetic scales, where there is often little 1:1 orthology of receptors among species (Tsutsui 2013; Yan et al. 2020). At the same time, studies of closely related species with similar ecologies and life histories can reveal the dynamics of receptor evolution at finer timescales and elucidate the process of gene family turnover (Guo & Kim 2007; Brand et al. 2015; Karpe et al. 2016; Brand & Ramirez 2017; Miller CH et al. 2020). Recent efforts to sequence a growing number of social insect genomes have suggested that social evolution is associated with expansions within the OR gene family, and the 9-exon OR subfamily in particular has experienced increased gene turnover and sequence evolution relative to other OR subfamilies (Zhou et al. 2012, 2015; Engsontia et al. 2015; Kapheim et al. 2015; Karpe et al. 2016, 2017; McKenzie et al. 2016). Given the importance of olfaction for social insect behavioral ecology, ORs provide a key route to linking genes to diverse and complex behaviors among ants, bees, and wasps. However, there are two major gaps in our knowledge of odorant receptor evolution in social insects. First, the hypothesis that social evolution is associated with OR expansions has yet to be tested in all of the independent origins of sociality among Hymenoptera. The independent origins of sociality in wasps provide an opportunity to compare patterns of OR gene family evolution to those that have been observed within social bees and ants (Hines et al. 2007). Second, studies of social insect ORs have focused on comparisons between genera or families, meaning the dynamics of OR gene family turnover across a gradient of species divergence times remain to be investigated. A better understanding of the short-term mechanisms of OR evolution provides additional insights into the molecular evolutionary dynamics shaping receptor diversity across more distantly related taxa. The recent release of five genomes of *Polistes* paper wasps spanning ~1-40 million years of divergence provide an opportunity to fill these gaps in our knowledge.

The molecular evolution of the OR gene family is best described as a birth-and-death process, in which genes are duplicated and deleted over evolutionary time (Nei 2007; Nozawa & Nei 2007; Eirín-López et al. 2012). Both random drift and natural selection are present during this process, determining the extent of OR gene copy number variation and the rate of gene sequence evolution (Nei 2007; Nozawa & Nei 2007). It is useful to subdivide odorant receptor function into behavioral and molecular functions, in order to examine how OR function at the organismal and molecular levels corresponds to different patterns of molecular evolution. Organisms use odorant receptor proteins to detect stimuli and provide input to neural circuits that regulate decision-making, here termed behavioral function (Yapici et al. 2014). At the molecular level, the specificity with which an OR binds chemical compounds or families of compounds (ligands) determines the tuning specificity of the olfactory receptor neuron (ORN) in which it is expressed (Hallem et al. 2004). ORN tuning specificity describes the relationship between the change in the frequency of action potentials fired by the ORN across a range of different chemical stimuli. An OR's molecular function can be categorized as specialist if ligand binding is highly specific, i.e. one chemical, or generalist if ligand binding allows for detection of multiple chemicals or chemical types. Narrowly tuned ORNs express specialist ORs that bind a specific ligand, creating a dedicated channel of olfaction, while broadly tuned ORNs express generalist ORs that bind a larger spectrum of ligands (Touhara & Vosshall 2009; Andersson et al. 2015). In some cases, ORNs achieve broad tuning by co-expressing multiple types of ORs (Fleischer et al. 2018). Generalist ORs are important for combinatorial coding: the process of combining input from multiple ORs that bind overlapping sets of ligands in order to discriminate a large variety of odors (Malnic et al. 1999). In nature, ORs exist on a spectrum from specialist to generalist, and ORN responses are dependent upon a complex interaction of OR, ligand concentration, and odorant binding proteins (Vogt et al. 1991; Hallem et al. 2004; Hallem & Carlson 2006; Stensmyr et al. 2012; Mathew et al. 2013; Dweck et al. 2015; Ebrahim et al. 2015; Münch & Galizia 2016).

There are distinct sets of predictions for the molecular evolution of ORs depending on their behavioral and molecular functions. Negative selection (purifying selection) is expected to conserve specialist ORs, which are not resilient to mutations due to the tight link between receptor and ligand in a dedicated olfactory channel (Andersson et al. 2015). For example, a volatile emitted by toxic microbes triggers avoidance behavior in *Drosophila melanogaster* by binding a specialist OR that is conserved across the genus (Stensmyr et al. 2012). In contrast to dedicated olfactory channels, combinatorial coding is thought to involve ORs with overlapping responses to ligands (Andersson et al. 2015). Mutations that slightly alter the response profiles of functionally redundant ORs may not be eliminated by negative selection, since other ORs can help compensate (Fishilevich et al. 2005; Keller & Vosshall 2007). Copy number variation and relaxed selection allow ORs to gain mutations that might endow them with an adaptive behavioral function. Divergent chemosensory landscapes between species lead to increased copy number variation as ORs with new behavioral functions are gained and ORs that detect irrelevant ligands are lost (Ramdya & Benton 2010; Goldman-Huertas et al. 2015).

The molecular evolution of the OR gene family is dynamic among the Hymenoptera, with prevalent lineage-specific gene expansions and losses, especially in the 9-exon OR subfamily (Engsontia et al. 2015; Zhou et al. 2015; McKenzie & Kronauer 2018). The 9-exon ORs constitute about one third of all ant ORs, and have evolved rapidly in ants, frequently under positive selection, leading researchers to propose that 9-exon ORs facilitate recognition of cuticular hydrocarbons (CHCs) (Smith CR, Smith CD et al. 2011; Smith CD, Zimin et al. 2011; Zhou et al. 2012, 2015; Engsontia et al. 2015; McKenzie et al. 2016). CHCs are used by insects to waterproof the cuticle and to communicate with conspecifics (Blomquist & Bagnères 2010). While less pronounced than in ants, dynamic evolution is also characteristic of 9-exon OR evolution in social bees, which rely on CHCs in communication (Sadd et al. 2015; Karpe et al. 2016, 2017). Functional studies in which ORs were transfected into an empty *D. melanogaster* ORN have verified that at least some 9-exon ORs of the ant *Harpegnathos saltator* overlap in their responses to ligands, with multiple 9-exon ORs responding to the same CHC molecule and unique 9-exon ORs responding to multiple different CHC molecules (Pask et al. 2017; Slone et al. 2017). Functional ORs are necessary for normal nesting behavior and for nestmate recognition in ants, a process which involves detecting variation in the CHCs on the cuticles of conspecifics (Lavine et al. 1990; van Zweden & d’Ettorre 2010; Sturgis & Gordon 2012; Trible et al. 2017; Yan et al. 2017; Ferguson et al. 2020). Together these studies suggest that 9-exon ORs function in combinatorial coding of CHC perception.

Like other Hymenopterans, vespid wasps, including the genus *Polistes*, use CHCs in complex social behaviors (Gamboa et al. 1986, 1996; Dani & Turillazzi 2018). *Polistes* use chemicals as signals and cues in a variety of behaviors, including during mate attraction, mate compatibility recognition, queen recognition, dominance/fertility signaling, and nestmate recognition (Reed & Landolt 1990; Espelie et al. 1994; Post & Jeanne 1984; Dapporto et al. 2007; Jandt et al. 2014; Sledge et al. 2001a, b, 2004; Oi et al. 2019). Recent efforts to sequence *Polistes* genomes provide an opportunity to resolve patterns of OR evolution among closely related species as an independent test of 9-exon OR gene subfamily expansion during social evolution (Patalano et al. 2015; Standage et al. 2016; Miller SE et al. 2020). We annotated the OR repertoires of five *Polistes* species representing ~40 million years of evolution: *P. fuscatus*, *P. metricus*, *P. dorsalis*, *P. canadensis*, and *P. dominula* (Figure 1). Combining neuroanatomy, manual gene annotation, and molecular evolution analysis, we examined the evolution of odorant receptors in light of the patterns predicted for different behavioral and molecular functions. We discover that social wasps, like ants, have an expanded subfamily of 9-exon ORs. Between *Polistes* species, 9-exon ORs exhibit dynamic evolution relative to ORs in other subfamilies, which are highly conserved. Expansion and birth-and-death evolution of the 9-exon OR subfamily in social wasps is consistent with a unique function in combinatorial coding perception of CHCs.

**Fig. 1:**
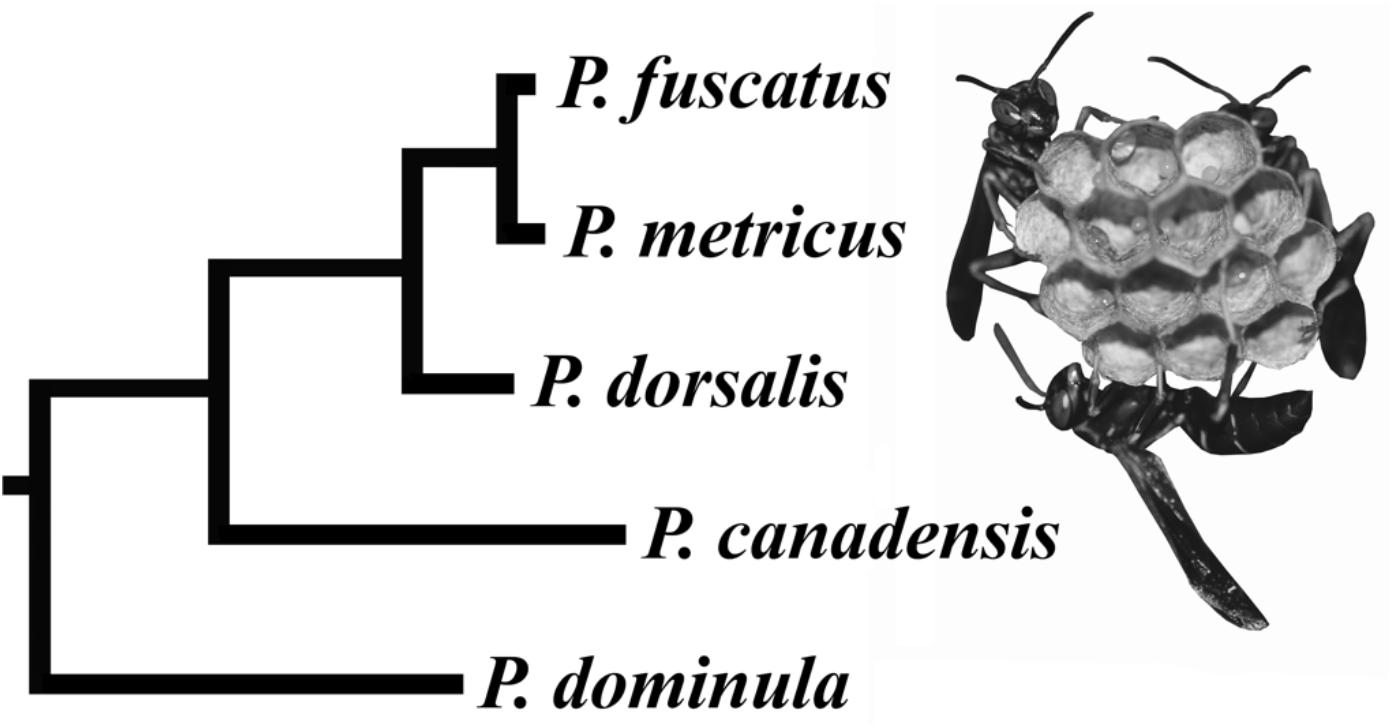
Phylogeny of five *Polistes* species considered in this study: *P. fuscatus*, *P. metricus*, *P. dorsalis*, *P. canadensis,* and *P. dominula*. The photo to the right of the phylogeny shows *P. fuscatus* foundresses on a nest. Phylogenetic tree based on the 16S ribosomal RNA gene and the cytochrome oxidase subunit I gene.

## RESULTS

### Antennal Lobe Neuroanatomy, Manual Gene Annotation Predict 200 ORs in *P. fuscatus*

In order to predict the OR repertoires of *P. fuscatus* and four other *Polistes* species, we combined fluorescent confocal microscopy of the *P. fuscatus* antennal lobe with manual genome annotation informed by antennal RNAseq. We found 229 glomeruli in the antennal lobe of an adult gyne (female reproductive) (Figure S1). Across a sample of insects, the number of intact OR genes in the genome correlates with the number of glomeruli in the antennal lobe, predicting 229 ORs in the *P. fuscatus* genome (Figure 2). Here we focus on the *P. fuscatus* genome because it has nearly chromosome level scaffolds and is the best assembled *Polistes* genome (Table S1; Patalano et al. 2015; Standage et al. 2016; Miller SE et al. 2020). Automated annotation using the MAKER pipeline (Holt & Yandell 2011) without guidance from antennal mRNA predicted 115 OR gene models in the *P. fuscatus* genome. A combined *P. fuscatus* male and gyne (reproductive female) antennal transcriptome generated using Trinity (Haas et al. 2013) yielded 89 OR genes greater than 900 nucleotides in length. Some long Trinity genes contain multiple 7-transmembrane domains and likely represent concatenated OR genes. The small fraction of the *P. fuscatus* OR repertoire predicted by transcriptome assembly is consistent with previous observations that annotation of OR repertoires using only transcriptome data typically fails to recover all ORs (Karpe et al. 2016, 2017, 2020).

**Fig. 2:**
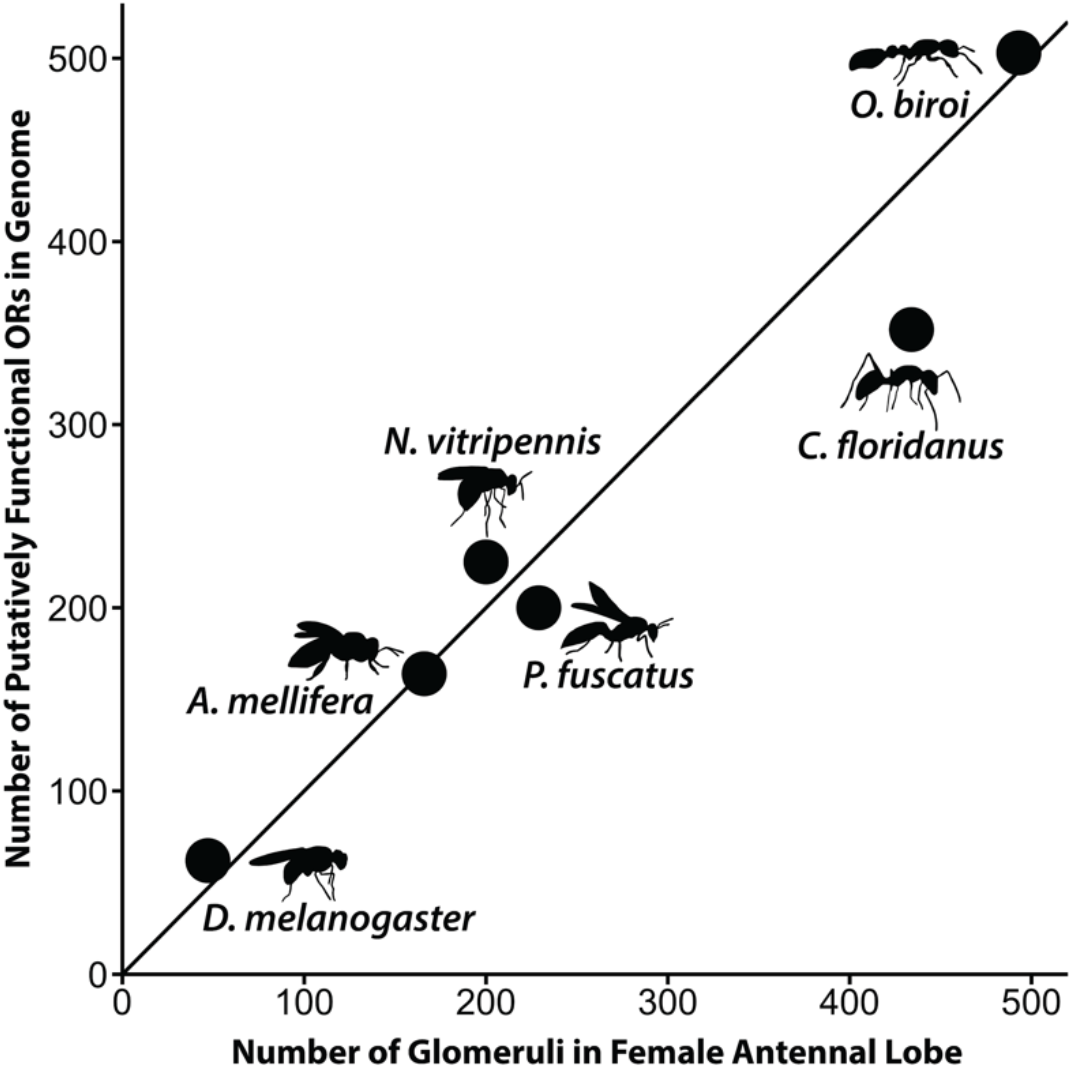
The number of functional ORs is correlated with the number of antennal lobe glomeruli across insect species (50 glomeruli and 62 ORs in the genome of the common fruit fly *D. melanogaster*, Fishilevich & Vosshall 2005; 166 glomeruli in the worker and 163 intact ORs in the genome of the honey bee *A. mellifera*, Arnold et al. 1985, Robertson & Wanner 2006; ~200 glomeruli in females and 225 intact ORs in the genome of the parasitic wasp *N. vitripennis*, Groothuis et al. 2019, Robertson et al. 2010; ~434 glomeruli in the worker and 352 functional ORs in the genome of the ant *C. floridanus*, Zube & Rössler 2008, Zhou et al. 2012; 493 glomeruli in the worker and 503 intact ORs in the genome of the ant *O. biroi*, McKenzie et al. 2016; McKenzie & Kronauer 2018). The diagonal line represents a line of equality with slope of 1.

Manual gene annotation of *P. fuscatus* ORs recovered 231 gene models across 28 scaffolds (Figure S2), of which 28 are pseudogenes and 10 are incomplete gene models (7 missing N termini, 2 missing C termini, and one missing both N and C termini). Since functional insect ORs are typically composed of 400 amino acids, we defined gene models as putatively functional if they coded for proteins greater than or equal to 300 amino acids in length. In *P. fuscatus*, the 200 putatively functional gene models encode protein sequences with an average length of 395 ± 15 (SD) amino acids, and 198 of these gene models encode protein sequences greater than 350 amino acids in length (Table 1). Odorant receptor proteins possess seven transmembrane domains (Wicher 2015). The putatively functional *P. fuscatus* OR proteins possess on average 5.95 ± 0.91 (SD) transmembrane domains as predicted by TMHMM version 2.0c (Sonnhammer et al. 1998) and 6.43 ± 1.13 (SD) as predicted by Phobius version 1.01 (Käll et al. 2004). For comparison, transmembrane domain prediction in 61 *D. melanogaster* ORs coding for proteins greater than 375 amino acids in length found on average 5.77 ± 1.12 (SD) transmembrane domains as predicted by TMHMM version 2.0c and 6.18 ± 1.09 (SD) as predicted by Phobius version 1.01 (sequences from Hopf et al. 2015 Supplemental Data 1). The close match between the number of ORs predicted by neuroanatomy and the number recovered from manual annotation suggests that we have identified nearly all of the OR genes in *P. fuscatus*. The number of transmembrane domains predicted are comparable to annotations of *D. melanogaster* and approach the 7 transmembrane domains expected for insect ORs. Manual OR gene annotation in *P. fuscatus* and four other *Polistes* genomes is summarized in Table 1.

**Table 1:**
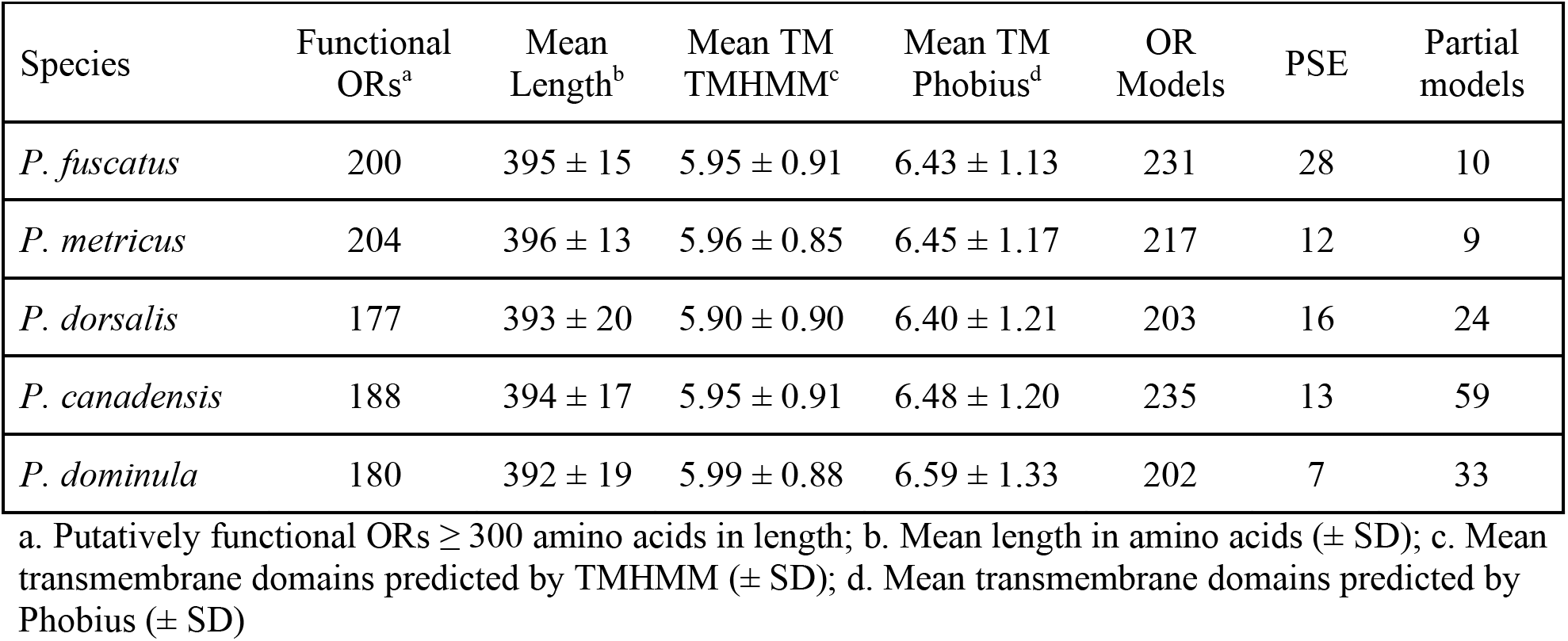
Summary of Odorant Receptor Gene Annotations in Five *Polistes* Genomes

### 9-exon OR Subfamily Expanded During the Evolution of Social Wasps

We conducted a Hymenoptera-wide analysis of OR evolution to test the prediction that the 9-exon OR subfamily was independently expanded during the evolution of eusociality in vespid wasps. By comparing the *P. fuscatus* OR repertoire to other Hymenopterans, our findings reinforce previous results showing that across Hymenopteran families, ORs evolve with lineage-specific expansions of multiple OR subfamilies (Figure 3). Gene gain and loss events were predicted using NOTUNG (Chen et al. 2000) and mapped onto a species cladogram of 14 Hymenopterans (Figure 4). NOTUNG estimated an ancestral Apocritan repertoire of 56 ORs, which has expanded independently during the evolution of braconid wasps, ants, bees, and paper wasps (Figure 4). The 9-exon subfamily is commonly expanded across Hymenoptera (~90 genes on average), and comprises ~36% of social insect OR repertoires. The largest lineage-specific expansions of Hymenopteran 9-exon ORs have occurred independently during the evolution of ants and social wasps. In *P. fuscatus*, this clade has expanded to 105 genes, comprising 53% of the OR gene set (Figure 4). Given the well-documented use of CHCs as signal molecules in *Polistes* (Singer 1998; Dani et al. 2001; Dani 2009), it is not surprising to find expansions in the CHC-detecting 9-exon subfamily in this genus. Subfamilies L, T, H, E, and V have also expanded in *Polistes*, but not to the extent of the 9-exon OR subfamily.

**Fig. 3:**
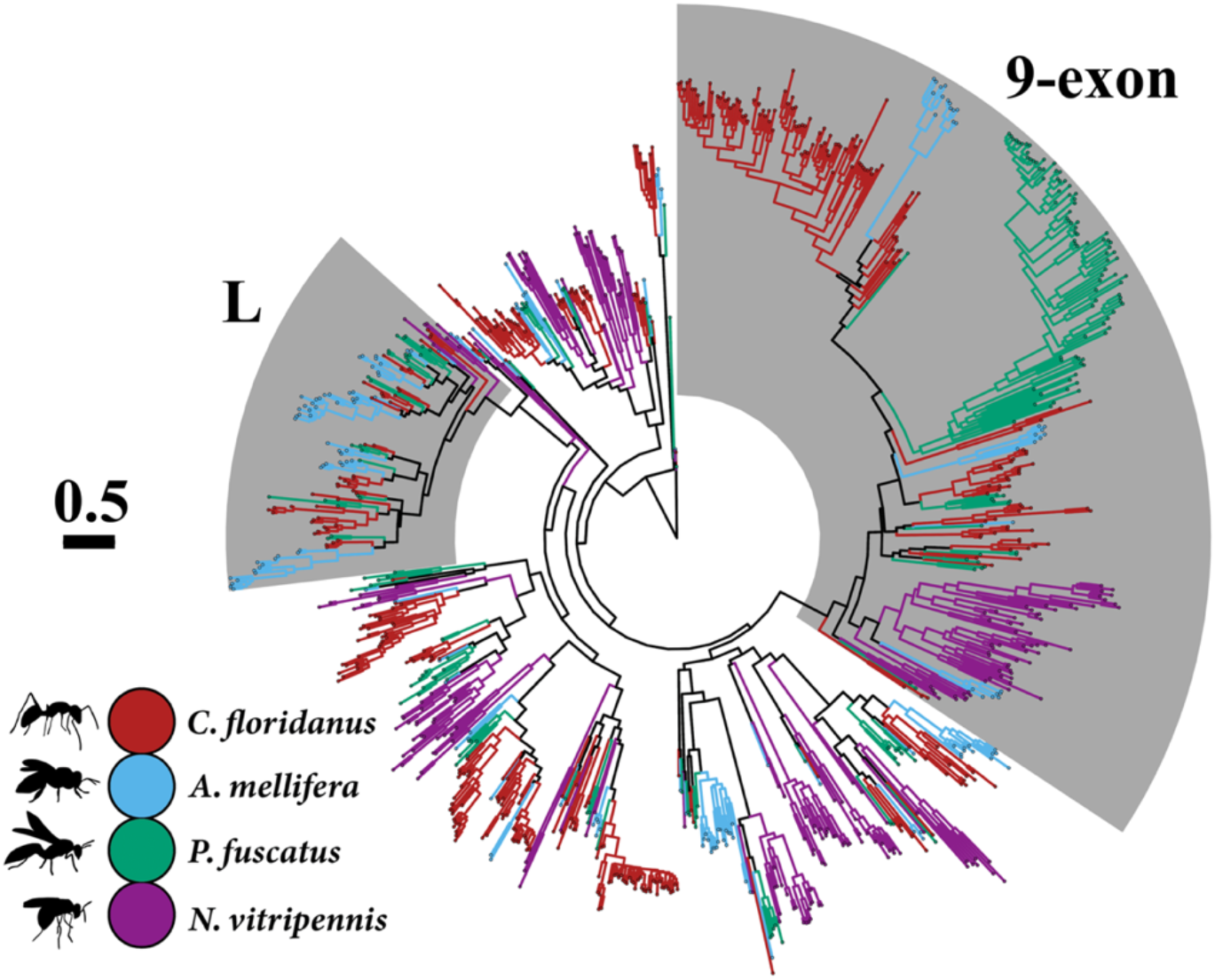
Maximum likelihood OR protein tree constructed using data from four Hymenopterans (*Apis mellifera*, Robertson & Wanner 2006; *Camponotus floridanus*, Zhou et al. 2012; *Nasonia vitripennis*, Robertson et al. 2010). Branches are colored by species (Red: *Camponotus floridanus*; Light blue: *A. mellifera*; Green: *P. fuscatus*; Purple: *N. vitripennis*). The L and 9-exon subfamilies are highlighted. Scale bar represents 0.5 mean substitutions per site.

**Fig. 4:**
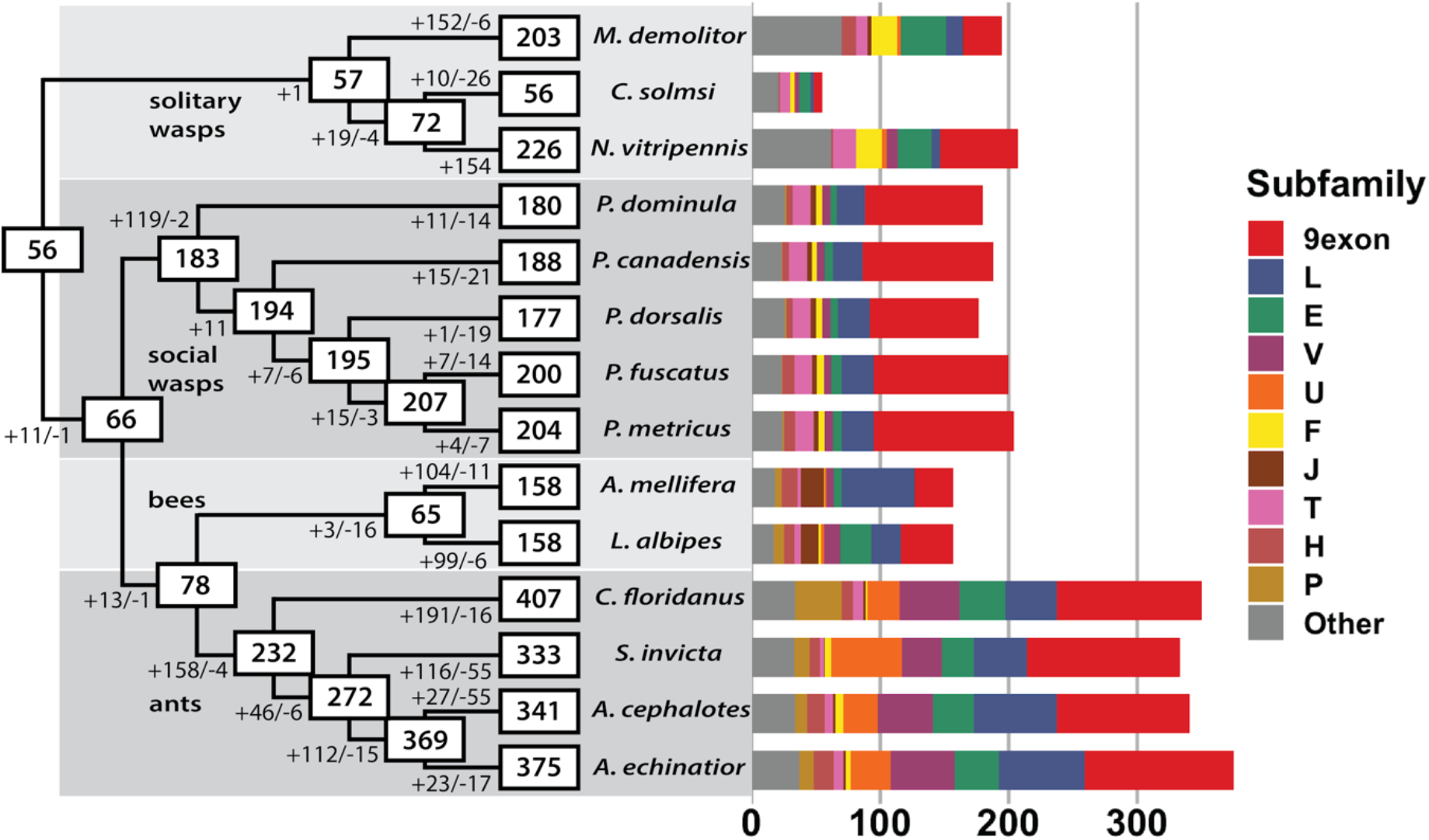
Cladogram of Hymenoptera species showing estimated number of OR gene gain and loss events along branches and estimated size of ancestral and extant species OR repertoires in boxes. To the right is a bar chart showing numbers of ORs broken down by subfamily. Non-Polistine OR data are from Robertson et al. 2010 and Zhou et al. 2012, 2015. The set of intact ORs that were longer than 300 amino acids was used except for *C. floridanus* in the bar chart, where only ORs considered putatively functional by Zhou et al. 2012 were used.

### The 9-exon OR Subfamily Shows a Distinct Pattern of Orthology Within *Polistes*

We next examined the evolutionary history of OR genes among the five *Polistes* species to reveal patterns of orthology and paralogy within subfamilies. Across the *Polistes* genus, most OR subfamilies are highly conserved (Figure 5A). About 70% of non-9-exon family *P. fuscatus* ORs are in 1:1 orthology with all other *Polistes* species sampled as predicted by OrthoFinder (Emms & Kelly 2015; Table S3). The remaining orthologous groups contain an expansion in one or more species (Figure 5B). Considering non-9-exon ORs, most ORs are shared by all five *Polistes* species examined, and most expansions are shared across all five species. Given that the species examined here span ~40 million years of divergence (Peters et al. 2017), the conservation of most of the OR repertoire is notable and may be related to the similarity of ecological and social niches found among *Polistes* wasps. While a common evolutionary history has led to large 9-exon OR complements in all *Polistes* species examined, lineage-specific gains and losses of 9-exon ORs account for most of the variation in OR repertoires size across *Polistes* species (Figure 4). In contrast with the other OR subfamilies, the 9-exon OR subfamily shows more lineage specificity with only 32% of *P. fuscatus* 9-exon ORs showing simple 1:1 orthology across all five *Polistes* examined (Table S3). Most 9-exon subfamily orthologous groups contain gene copies from four or fewer species, and lineage-specific expansions are more common in 9-exon OR orthologous groups (Figure 5B). The relative lack of orthology among 9-exon OR genes compared to the rest of the OR gene subfamilies suggests unique evolutionary processes shaping 9-exon ORs.

**Fig. 5:**
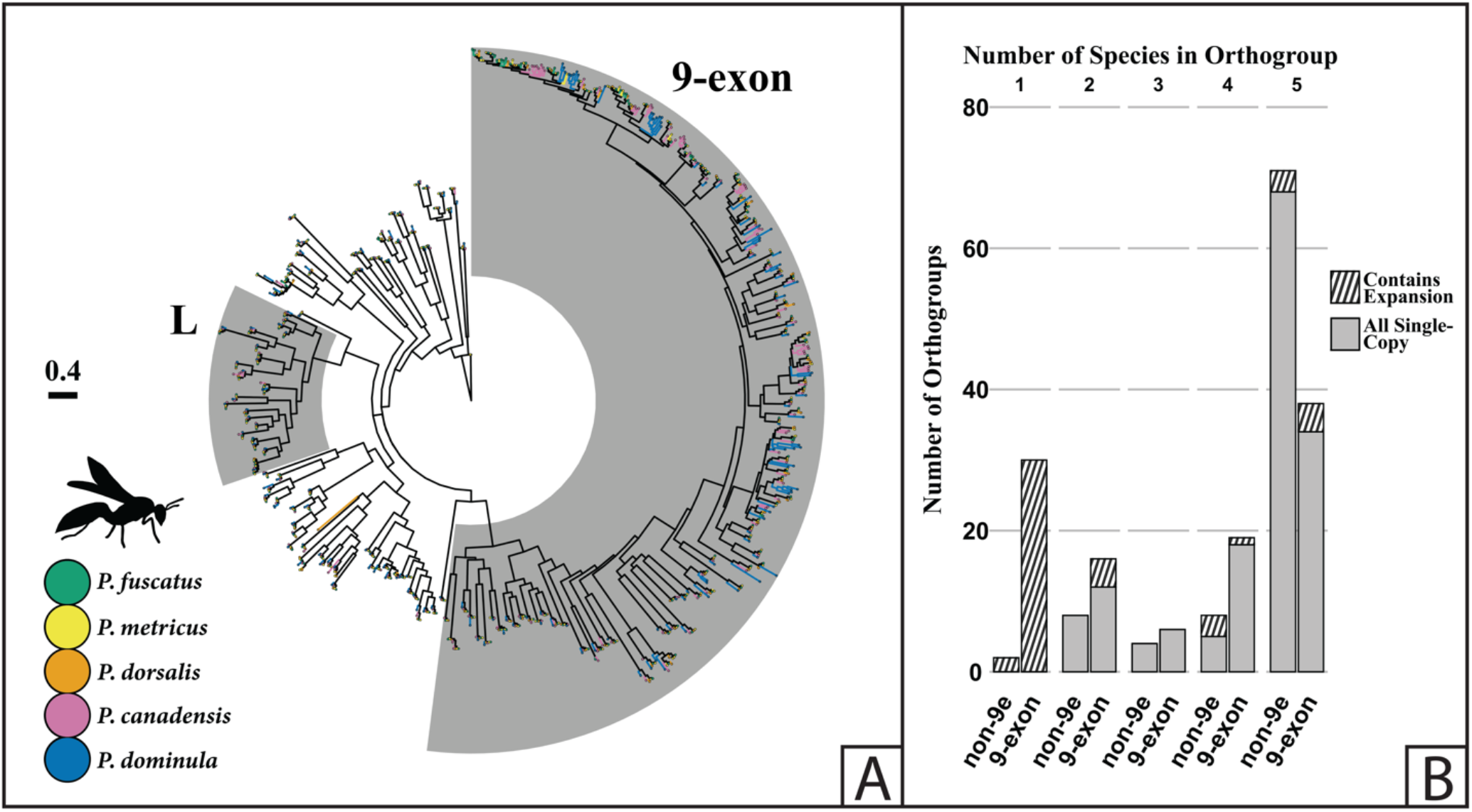
(A) Maximum likelihood OR protein tree with branches colored by species (Green: *P. fuscatus*; Yellow: *P. metricus*; Orange: *P. dorsalis*; Magenta: *P. canadensis*; Blue: *P. dominula*). The L and 9-exon subfamilies are highlighted. Scale bar represents 0.4 mean substitutions per site. (B): Stacked bar chart showing the number of *Polistes* species (x-axis) represented in each orthologous group (y-axis), and whether or not each orthologous group is single copy (shaded bottom portion of bar) or contains an expansion in at least one species (top striped portion of bar). Orthologous groups are split into two categories: non-9-exon orthologous groups (left bar) and 9-exon orthologous groups (right bar).

### Microsynteny Reveals Recent Birth-and-Death Events in *Polistes* 9-exon OR Subfamily

Expanded gene families often occur as tandem arrays, a genomic architecture that can contribute to increased rates of gene birth and death, increasing copy number variation among species (Ohno 1970). Therefore, we examined how genomic organization varies between OR subfamilies in *Polistes* species to generate insights into the molecular evolutionary mechanisms shaping OR subfamily function. Genomic organization of ORs across *Polistes* is consistent with a model of birth-and-death evolution shaping OR repertoires. As in bees, gene gain and loss at a small number of loci containing tandem arrays is responsible for most copy number variation in the OR family across closely-related species (Brand & Ramírez 2017). In *P. fuscatus*, 62% of ORs occur in tandem arrays of 6 or more genes (Figure 6A). The frequency of tandem arrays and the tail-to-head orientations of neighboring genes point to tandem duplication as the primary mechanism of OR expansion, likely caused by non-allelic homologous recombination (Lynch 2007; Ramdya & Benton 2010). We examined microsynteny among genes and pseudogenes in the four longest tandem arrays of ORs in *Polistes* genomes (Figure 6B). There is marked decrease in OR synteny among genes in orthologous 9-exon OR arrays compared to tandem arrays of L and T subfamily ORs in the *Polistes* genus. The longest OR gene tandem array in *P. fuscatus* is comprised of 44 genes in the 9-exon subfamily on scaffold 13 (*s13*), which corresponds to homologous arrays of 50 genes in *P. metricus*, 25 genes in *P. dorsalis*, 33 genes in *P. canadensis*, and 29 genes in *P. dominula*. Only 34% of *P. fuscatus* ORs in this array have orthologs across all *Polistes* species sampled (Figure 6B). The second longest OR gene tandem array in *P. fuscatus* contains 24 ORs in the L subfamily on scaffold 17 (*s17*), and these ORs show 1:1 orthology across *P. fuscatus*, *P. metricus*, and *P. dorsalis*, while *P. canadensis* possesses an array of ~23 genes split between two scaffolds, and *P. dominula* possesses an array of 21 ORs at this locus (Figure 6B). This tandem array, widely expanded across Hymenoptera, has been expanded and conserved across *Polistes*. The T subfamily, located on scaffold 8 (*s8*) of the *P. fuscatus* genome, is composed of 14 tandemly arrayed genes that show 1:1 orthology across five *Polistes* (Figure 6B). Differences in the extent of microsynteny among tandem arrays belonging to different OR subfamilies highlight the unique evolutionary processes shaping 9-exon OR evolution in paper wasps. At the same time, the extreme conservation of L and T subfamily tandem arrays across species highlights the strong conservation of the OR repertoire outside of the 9-exon subfamily among *Polistes* species.

**Fig. 6:**
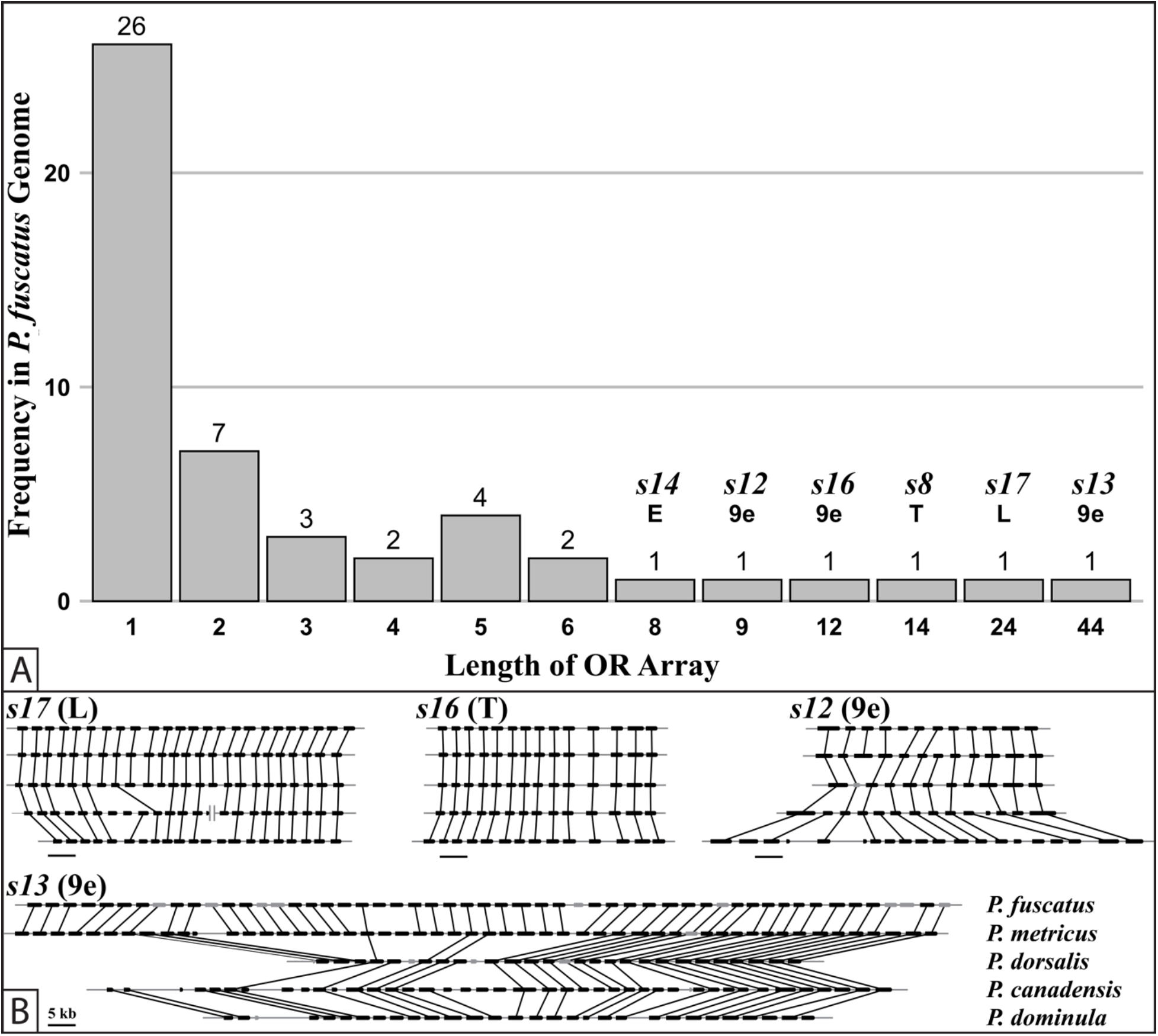
(A) Frequency of OR gene singletons and tandem arrays in the *P. fuscatus* genome. 62% of ORs in *P. fuscatus* occur in tandem arrays of 6 or more genes. The longest tandem array is a 44 gene cluster on scaffold 13 (*s13*) containing 9-exon subfamily ORs. The first row of x-axis labels is the number of OR genes in a tandem array cluster, and the second row labels the OR subfamily and scaffold number (abbreviated *s#* in parentheses) of the six longest tandem arrays. (B) Genome alignments of four loci containing tandem arrays of OR genes in all *Polistes* species examined. Each alignment is labeled with the corresponding OR subfamily and *P. fuscatus* scaffold number (abbreviated *s#* in parentheses). Black boxes represent presumably functional genes and gray boxes represent pseudogenes. Directionality of genes is denoted by curved corners at the 3' (tail) end. Black lines connect orthologous genes between species. Genomic scaffolds are represented by horizontal, gray lines, and scaffold ends are represented by vertical gray lines. The black scale bars represent 5 kb.

Microsynteny analysis suggests a process of ongoing gene turnover in 9-exon arrays but stasis in most other expanded subfamilies. More recent turnover should be associated with higher pairwise amino acid identity between neighboring genes in an array if they are the result of recent duplication events (Ohno 1970; Bohbot et al. 2007). To explore the relationship between amino acid divergence and tandem array locus, we compared the mean percent amino acid identity among neighboring genes within an array between the eight loci containing the longest tandem arrays of ORs in the *P. fuscatus* genome using one-way ANOVA (Figure 7). Mean percent amino acid identity of neighboring genes was significantly separated by OR array identity (DF = 7; F = 5.39; P = 2.67e-05). Differences between particular OR tandem arrays were identified using Tukey HSD post hoc tests. The mean percent amino acid identity among neighboring genes within one tandem array of nine 9-exon ORs on scaffold 12 (*s12*) of the *P. fuscatus* genome is higher than in the *s13* 9-exon array (P Adj = 0.04586), the *s17* L array (P Adj = 0.00013), the s8 T array (P Adj = 0.00458), the *s16* 9-exon array (P Adj = 0.00124), and the *s19* V array (P Adj = 0.02247). The *s12* 9-exon OR array is composed of a larger proportion of pseudogenes (5 PSE, 10 intact gene models) than the other two 9-exon arrays (*s13*: 9 PSE, 44 intact gene models; *s16*: 0 PSE, 12 intact gene models). ORs in the *s12* 9-exon array lack clear orthologous relationships with ORs in species other than *P. metricus*. Taken together, the high within array sequence similarity, high frequency of pseudogenes, and low orthology exhibited by this array indicate that it is the result of one or more recent gene duplication events since the divergence of *P. fuscatus* and *P. metricus* from the other three *Polistes* species. The *s6* H subfamily array also shows higher amino acid sequence identity among neighboring genes than the *s17* L subfamily array (P Adj = 0.01625) and the *s16* 9-exon array (P Adj = 0.04038). Increased amino acid similarity may also occur within older tandem arrays as a result of gene conversion (Nagawa et al. 2002). However, we searched for gene conversion using GENECONV (Sawyer 1989) and did not detect gene conversion events within the *s12* 9-exon array or in the *s6* H array after Bonferroni correction. Patterns of genomic organization of OR genes in *Polistes* genomes lead to the conclusion that gene gain and loss in the 9-exon OR subfamily is an ongoing process within this genus, in contrast to the stable and conserved tandem arrays in most other OR subfamilies.

**Fig. 7:**
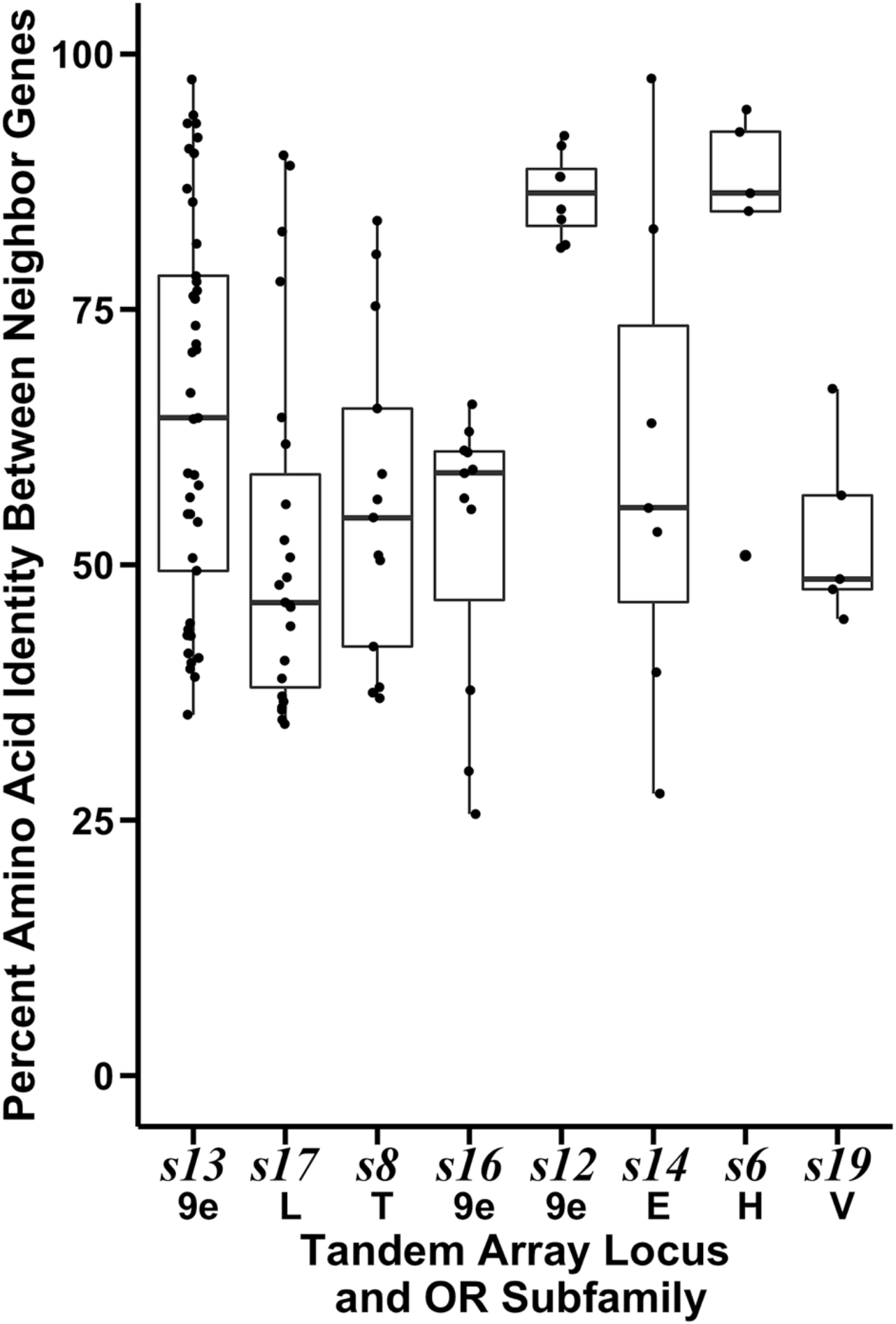
Percent amino acid identity between neighboring genes at eight loci containing the longest OR gene tandem arrays in the *P. fuscatus* genome. Arrays are ordered by length in gene number, from longest (44 9-exon subfamily ORs in the *s13* tandem array) to shortest (6 H subfamily ORs in the *s6* tandem array and 6 V subfamily ORs in the *s19* tandem array).

### Positive Selection in Expanded OR Subfamilies and Accelerated Evolution of 9-exon ORs

We examined patterns of OR evolution among five *Polistes* species to test the prediction that OR subfamilies exhibit signatures of positive selection. We were especially interested in whether the recent dynamism in 9-exon OR gene copy number has been accompanied by episodes of positive selection in *Polistes*. HyPhy aBSREL (Smith MD et al. 2015; Pond et al. 2020) analyses of *Polistes* OR subfamilies detected eight branches under episodic positive diversifying selection, all in OR subfamilies with expansions: three branches in the 9-exon subfamily (0.33% of 918 9-exon subfamily branches; Figure S4); three branches in the L subfamily (1.28% of 234 L subfamily branches; Figure S5); one branch in the E subfamily (1.67% of 60 E subfamily branches; Figure S6); and one branch in the H subfamily (1.54% of 65 H subfamily branches; Figure S7). This supports the hypothesis that gene duplication releases duplicate genes from selective constraints, allowing duplicate sequences to evolve towards other evolutionary optima (Ohno 1970). While the 9-exon OR subfamily is not unique among expanded OR subfamilies in its instances of episodic positive selection as measured by HyPhy aBSREL, the rate of amino acid divergence is higher among the 9-exon OR subfamily as a whole. The 91% mean amino acid identity among 1:1 9-exon subfamily orthologs in *Polistes* is significantly lower than the 95% mean amino acid identity among 1:1 orthologs in all other OR subfamilies (Figure S8; Welch Two Sample t-test: P = 7.051e-05).

To further evaluate the patterns of nucleotide substitution driving accelerated amino acid evolution of 9-exon ORs, we computed the values of *d*_N_ and *d*_S_ for pairwise alignments of 150 single copy orthologs between *P. fuscatus* and *P. dorsalis* (Figure 8) using model yn00 of PAML (Yang 2007). Values of *d*_N_ are significantly higher in 9-exon (mean *d*_N_ = 0.015) compared to other OR ortholog pairs (mean *d*N = 0.006) (Welch Two Sample t-test, P = 5.317e-07). Values of *d*_S_ are not significantly elevated among 9-exon ortholog pairs compared to other OR subfamilies (mean *d*_S_ = 0.029 in 9-exon ORs and 0.025 in non-9exon ORs, P = 0.343). Omega values (*d*N/*d*_S_) greater than 1 are often considered evidence of positive selection, while *d*N/*d*_S_ = 1 corresponds to neutral drift, and *d*N/*d*_S_ < 1 is evidence of negative selection. The omega value (*d*N/*d*_S_) for all genes is less than one, suggesting negative selection. However, omega is significantly higher in 9-exon ORs than in non-9-exon ORs, indicating that negative selection is weaker on 9-exon ORs (Welch Two Sample t-test: mean omega = 0.32 in non-9-exon ORs and 0.644 in 9-exon ORs, P = 8.027e-05). In general, negative selection conserves ORs shared by *P. fuscatus* and *P. dorsalis* (mean omega = 0.454), but an elevated rate of non-synonymous substitutions in 9-exon ortholog pairs imply relaxed negative selection and more drift responsible for sequence evolution in the 9-exon relative to other OR subfamilies.

**Fig. 8:**
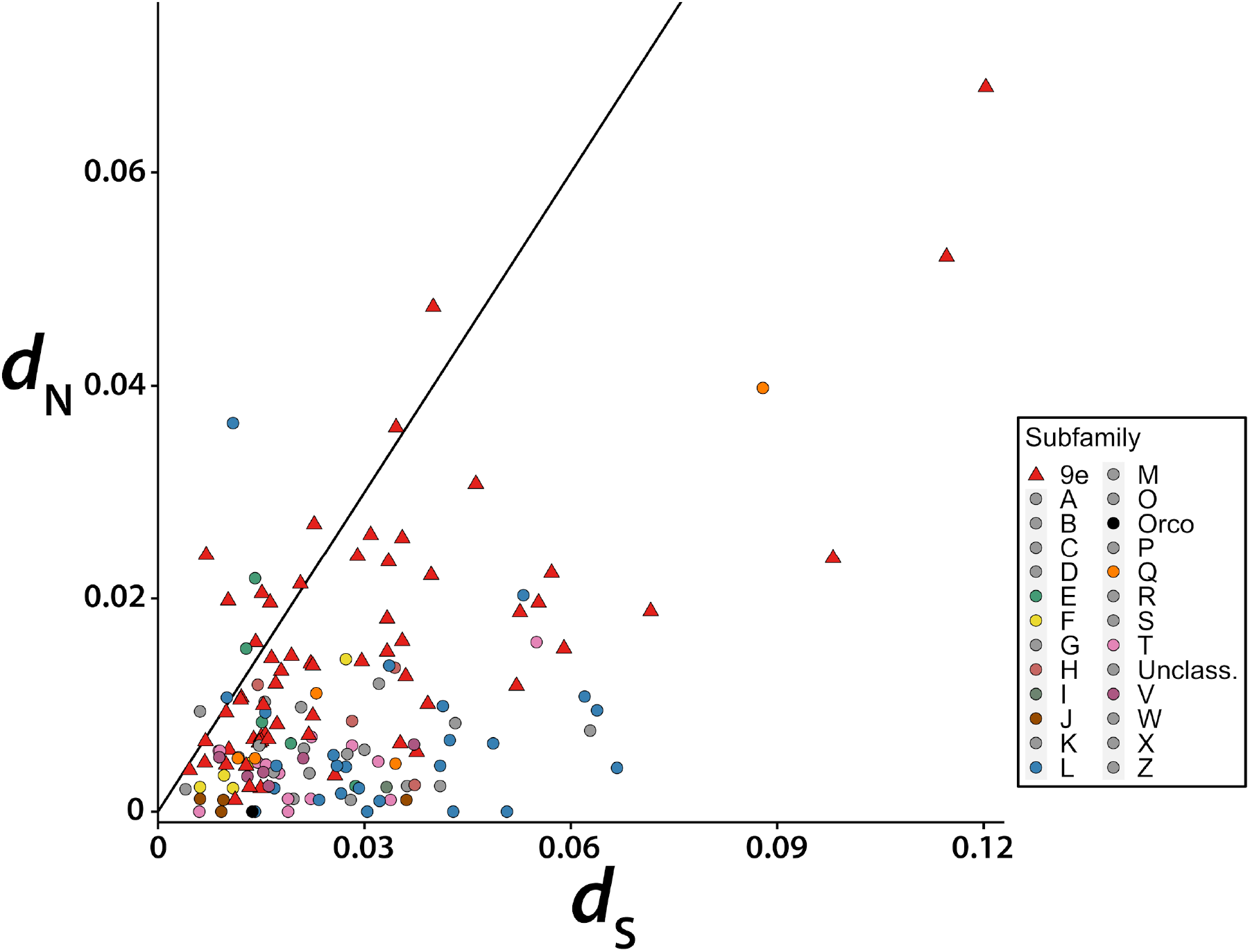
The values of *d*_S_ (x-axis) and *d*_N_ (y-axis) from pairwise alignments of *P. fuscatus* and *P. dorsalis* 1:1 orthologs. Values of *d*_N_ are elevated in the 9-exon OR subfamily (data points represented by red 393 triangles) relative to other OR subfamilies (data points represented by circles). The diagonal line represents a line of equality with slope of 1.

## DISCUSSION

### Expansion of 9-exon OR Subfamily During Independent Evolution of Sociality in Wasps

By carefully annotating the OR repertoires of five social wasp species spanning ~40 million years of divergence in the *Polistes* genus, this study adds a higher resolution lens to our view of the evolution of social insect odorant receptors. During the diversification of *Polistes,* evolutionary patterns show genus-wide conservation of their ~200 ORs except for the 9-exon genes, which show elevated turnover and lower sequence conservation. The 9-exon OR subfamily has dramatically expanded in paper wasps, and now makes up over half of the *Polistes* OR gene set. Social and ecological niches are relatively conserved within *Polistes*, though there is considerable variation in social behavior and ecological niches among vespid wasps (Ross & Matthews 1991; O'Neill 2001). For example, an analysis of three new, high-quality hornet genomes suggested that the highly eusocial hornets have larger OR repertoires compared to the primitively eusocial *Polistes* (Harrop et al. 2020). That analysis recovered less than half of the ORs reported here for *Polistes*, likely due to a lack of antennal transcriptome data, suggesting that hornets may have larger OR repertoires than reported. Evidence from the hornet *Vespa velutina*, including the discovery of 264 antennal lobe glomeruli, indicates that the hornet OR repertoire has expanded (Couto et al. 2016, 2017). Future analysis of additional genomes and antennal transcriptomes of diverse social and solitary vespid wasps will allow further examination of the relationship between social behavior and OR subfamily expansion.

### Combinatorial Coding of CHCs by 9-exon ORs Facilitates Recognition

Electrophysiological deorphanization studies of 9-exon ORs in the ant *Harpegnathos saltator* offer key insights into how 9-exon OR coding might relate to gene expansion. Through combinatorial coding, 9-exon ORs can detect a large variety of structurally diverse CHCs. Pask et al. (2017) examined 22 *H. saltator* 9-exon ORs, a subset of the 118 annotated 9-exon ORs in this species, and found that 9-exon ORs were responsive to CHCs, and overlapped in their responses to multiple CHC compounds. The combined responses of these 22 ORs to CHC extracts from different castes were sufficient to map the CHC profiles of males, workers, and reproductive females (gamergates) to separate regions of a 22-dimensional receptor space (Pask et al. 2017). This highlights the ability of 9-exon ORs to facilitate social recognition by combinatorial coding. In social insect colonies, CHC variation holds information at multiple levels of conspecific recognition, from inter-colony nestmate recognition to within colony individual recognition (d'Ettorre & Moore 2008; Leonhardt et al. 2016). Expansion of the 9-exon OR subfamily might result from selection for more combinations of ORs that together can discriminate between subtle qualitative and quantitative variations in CHC blends of conspecifics. Nest-specific quantitative variation in CHCs has been documented across *Polistes* species (Espelie et al. 1990; Singer et al. 1992; Espelie et al. 1994; Layton et al. 1994), but the molecular mechanisms underlying nestmate recognition in *Polistes* are still obscure. Increased copy number of 9-exon ORs may not only expand the qualitative range of compounds perceived by paper wasps, but also the quantitative olfactory space, since wasps may be able to discern unique concentration differences between CHC blends as a result of the combined action of 9-exon ORs with various response thresholds. Gene duplication can also promote regulatory diversification (Kucharski et al. 2016; Dyson & Goodisman 2020). In *P. metricus*, CHCs vary between castes and across stages of the colony cycle (Toth et al. 2014). Regulatory subfunctionalization of duplicate ORs could be responsible for caste- and colony phase-specific expression of ORs involved in detecting caste-specific and seasonally variable CHCs. In addition to adaptive expansion of ORs, neutral processes contribute to OR gene birth-and-death events. There may be an advantage for a large 9-exon OR gene copy number up to a point, followed by random gene duplication and deletion around this optimal copy number. This random genomic drift has been proposed to shape vertebrate olfactory receptor evolution and copy number variation in other large multigene families (Nei 2007; but see Hayden et al. 2010).

### Evolution of Odorant Receptors Reflects Specific Chemosensory Ecologies of Species

Social insect species differ in their level of sociality and extent of olfactory recognition abilities (d'Ettorre & Moore 2008; Rehan & Toth 2015). Some social aspects of the *Polistes* colony cycle vary across species. For example, the average number of cooperative foundresses varies from 1 to ~6, and average sizes of mature nests may vary ~60 to ~490 cells (Reeve 1991; Sheehan et al. 2015; Miller SE et al. 2018). Increased 9-exon OR copy number may facilitate complex olfactory recognition in species with larger colony sizes, higher cooperative nest-founding rates, and greater sympatry with related species. However, expansions of 9-exon ORs are not exclusive to social wasps, suggesting that the specific chemical ecology of an insect is a more influential factor shaping OR evolution than level of sociality (Karpe et al. 2017). Furthermore, a meta-analysis found that the complexity of CHC phenotypes does not differ between social and solitary Hymenopteran species (Kather & Martin 2015). The CHC profile of *Nasonia vitripennis* includes at least 52 CHC compounds, and detection of CHCs on prey items may help *Microplitis* identify prey (Lewis et al. 1988; Niehuis et al. 2011). The need for solitary wasps to perceive CHCs could explain why *N. vitripennis* and *M. demolitor* exhibit expansions in the 9-exon OR subfamily.

### Lineage-Specific Chemical Signaling in *Polistes* and Molecular Evolution of ORs

Most expanded OR subfamilies are highly conserved in copy number across five *Polistes* species, with the exception of the 9-exon OR subfamily. In particular, one portion of the 9-exon subfamily arranged in a single tandem array (*P. fuscatus* 9e *s13*) has experienced dynamic evolution. What might drive rapid gain and loss of 9-exon ORs? Divergent social chemical landscapes between species may cause gene turnover as 9-exon OR evolution tracks evolutionarily labile chemical signals. For example, *P. fuscatus* and *P. metricus* are closely related, and both species possess CHC profiles consisting of linear and methyl-branched alkanes (Espelie et al. 1990; Espelie et al. 1994). However, the *P. fuscatus* CHC profile includes a higher proportion of alkenes than *P. metricus* or *P. dominulus*, and the position of the methylated carbon of methyl-branched alkanes is sometimes shifted between species (Espelie et al. 1990; Singer et al. 1992; Espelie et al. 1994; Layton et al. 1994). Ant 9-exon ORs respond differently to subtle variations in CHC structure (Pask et al. 2017). Between closely related *Polistes* species, structural isomers of methyl-branched alkanes probably activate different ensembles of ORs.

If a chemical develops new behavioral relevance in a lineage, gene duplication would allow the olfactory system to explore chemical space in the direction of this compound. HyPhy aBSREL analyses identified eight branches in expanded OR subfamilies, including the 9-exon subfamily, that have undergone positive selection during the last ~40 million years, consistent with neofunctionalization or subfunctionalization of duplicated genes. Signatures of positive selection on OR genes may indicate directional selection to perceive species-specific chemical signals. Perception of such species-specific CHCs might be important in mate compatibility recognition. In *Polistes*, mating occurs at territories defended by males and often frequented by multiple species (Post & Jeanne 1983; Reed & Landolt 1990). However, the frequency of interspecific mating is low, suggesting *Polistes* use vision and/or olfaction to inform their mating decisions (Miller SE et al. 2019). Duplication and deletion of ORs would facilitate evolution of species-specific chemical signaling systems that could contribute to reproductive isolation of sympatric species. If a chemical signal is lost in a species, the corresponding ORs may become obsolete, and would be expected to pseudogenize and be purged from the genome. Duplication and deletion of ORs could also lead to species-specific chemical signaling in the absence of evolutionary change in chemical signals (Cande et al. 2013). However, OR evolution is not strictly necessary for such a difference to evolve between species, and circuit-level changes can prescribe new valence to chemical signals that are shared between species and perceived by common peripheral receptors (Seeholzer et al. 2018).

### Conservation of Most OR Subfamilies Suggests Conserved Functions

Aside from the 9-exon OR subfamily, gene expansions have occurred in subfamilies L, T, H, E, and V (Figure 4). A larger variety of ORs relaying information through ORNs to a larger number of antennal lobe glomeruli will increase sensory acuity in any olfactory discrimination task, social or otherwise. An ancient locus of tandemly duplicated L subfamily ORs observed across social insects has expanded in *Polistes*, although to a lesser extent than in other social insects (~50 L subfamily ORs in honeybee and ants, 25 L subfamily ORs in a tandem array on *P. fuscatus* scaffold 17). Odorant receptors in the L subfamily are thought to detect queen pheromone components and fatty acids in bees as well as CHCs in ants (Wanner et al. 2007; Karpe et al. 2016; Pask et al. 2017). The T subfamily has expanded to a greater degree in *P. fuscatus* (14 genes) than in ants (~7 genes) and the honeybee (2 genes), but no ORs in this clade have been functionally characterized. The *P. fuscatus* genome encodes nine H subfamily ORs, which are putative floral odorant detectors in bees, and which also respond to CHCs and other general odorants in ants (Claudianos et al. 2014; Slone et al. 2017). Fatty acids and volatile organic compounds are produced by flowers that wasps rely on as a source of carbohydrates (Raguso 2008). Expansions in several OR subfamilies may increase olfactory discrimination of chemicals with diverse behavioral relevance. *Polistes* species are distributed globally in temperate and tropical regions, occupying similar social and ecological niches as generalist predators and floral foragers that form primitively eusocial societies (Reeve 1991; Richter 2000). A conserved set of ORs may perform common functions in conserved behaviors across paper wasp species. High levels of OR conservation are also consistent with a specialist molecular function of an OR in a dedicated channel of olfaction. Patterns of molecular evolution suggest conserved behavioral and molecular functions of most non-9-exon OR subfamilies in *Polistes*.

### Distinct Patterns of OR Evolution Within the Same Genome

The differences between the conserved OR repertoires in *Drosophila* and the more dynamic evolution of vertebrate OR gene families have given rise to speculation about the relationship between OR function and evolution (Nozawa & Nei 2007; Andersson et al. 2015). There is prevalent negative selection conserving odorant receptors across *Drosophila* species, and the majority of *D. melanogaster* ORs form simple orthologous relationships across the genus (Clark et al. 2007; Guo & Kim 2007; McBride & Arguello 2007; Nozawa & Nei 2007; Sánchez-Gracia et al. 2009; Mansourian & Stensmyr 2015). In paper wasps we report both highly conserved OR expansions similar to those seen in *Drosophila* as well as elevated gene turnover and drift among the 9-exon ORs, reminiscent of a more vertebrate-like evolutionary pattern. If the highly dynamic clades of 9-exon ORs of social wasps are involved in more combinatorial coding compared to other more conserved 9-exon or non-9-exon ORs, that would indicate a link between molecular evolution of odorant receptors and neural coding. Further investigations into the relative tuning of 9-exon as well as more conserved ORs in social wasps and other social insects provide a promising research direction to investigate the links between molecular evolutionary patterns, odorant receptor tuning, and neural coding.

## MATERIALS & METHODS

### Antennal Lobe Imaging

Antennal lobe glomeruli of male and female *P. fuscatus* wasps were stained with anti-synapsin and imaged using a confocal laser scanning microscope. Details of the immunocytochemistry and imaging are included as Supplementary Materials & Methods.

### Gene Annotation

The *P. fuscatus*, *P. dorsalis*, and *P. metricus* genomes were assembled and automatedly annotated as described in Miller SE et al. (2020). The *P. canadensis* and *P. dominula* genomes and annotations were accessed through NCBI (Patalano et al. 2015; Standage et al. 2016). Coding regions of ORs were identified by using TBLASTN (Altschul et al. 1997) with a sample of OR proteins from 19 insect species used as query sequences. Genomes were queried iteratively with TBLASTN, adding newly annotated *Polistes* ORs to the query file, until no new OR coding regions were identified. To guide annotation of exon-intron boundaries, antennal mRNA from *P. fuscatus* males and females (gynes) was mapped to *P. fuscatus*, *P. metricus*, and *P. dorsalis* genomes using STAR (Dobin et al. 2013) and assembled into transcripts using Trinity (Table S2; Haas et al. 2013). Predicted transcripts were aligned to genomes using BLAT. Uncertain gene models in *P. metricus*, *P. dorsalis*, *P. canadensis*, and *P. dominula* were aligned to their orthologs in *P. fuscatus* using Muscle version 3.8.425 with maximum 4 iterations (Edgar 2004), and gene models were manually adjusted. All annotation evidence was imported into Geneious v11.1.5 genome browser for manual annotation. The majority of apparently functional ORs that were not detected by the automated annotation and required extensive manual curation were 9-exon subfamily receptors (e.g. Figure S9). Gene models were called pseudogenes if they exhibited frame-shift mutation, premature stop codons, or unacceptable 5’ donor or 3’ receptor splice sites. Transmembrane helices of all putatively functional ORs (>300 amino acids) were predicted using TMHMM version 2.0c (Sonnhammer et al. 1998) and Phobius version 1.01 (Käll et al. 2004). Throughout the main text, “putatively functional ORs” are OR proteins at least 300 amino acids in length. Details of mRNA library preparation, sequencing, read mapping, and manual gene annotation are included as Supplementary Materials & Methods.

### Phylogenetic Reconstruction

Phylogenetic trees were constructed using RAxML (Stamatakis 2014, Jones et al. 1992). Gene duplication and loss events were reconstructed by reconciling a gene tree with a species tree in NOTUNG version 2.9.1.3 (Durand et al. 2006; Vernot et al. 2008). Orthologous genes were determined using OrthoFinder (Emms & Kelly 2015), bootstrap support, and microsynteny. Phylogenetic reconstruction methods are explained in detail in the Supplementary Materials & Methods.

### Genomic Organization

OR genes and pseudogenes were considered to be in a tandem array if they were uninterrupted by non-OR genes and were within 5kb of each other. The lengths of OR arrays correspond to the number of putatively functional ORs and exclude the pseudogenes contained within the array. The pairwise percent amino acid identity between neighbors in an array was calculated using only putatively functional ORs that neighbored another putatively functional OR within 5kb. See Supplementary Materials & Methods for details.

### Sequence Analyses

All putatively functional *Polistes* ORs greater than 350 amino acids in length were used in analyses of episodic diversifying positive selection and gene conversion within OR subfamilies using aBSREL in HyPhy version 2.5.15 (Smith MD et al. 2015; Pond et al. 2020) and GENECONV (Sawyer 1989). Values of pairwise *d*N/*d*_S_ for orthologs shared by *P. fuscatus* and *P. dorsalis* were calculated using PAML version 4.9 (Yang 2007) program yn00 with the Yang & Nielsen (2000) method. See Supplementary Materials & Methods for details.

## Supporting information

Polistes ORs fasta, gff, and subfamily CDS alignment

Supplementary table containing OR metadata

## Data Availability

The genome assemblies analyzed in this article are available on Genbank (see Table S1). Gene models, amino acid sequences, and nucleotide sequences underlying this article, as well as alignments analyzed in selection analyses, are available in its online Supplementary Material.

## SUPPLEMENTARY MATERIAL

Supplementary Materials & Methods, Supplementary Tables S1-S3, and Supplementary Figures S1-S9 are included in a PDF file available in the online Supplementary Material. Meta-data associated with *Polistes* ORs, including subfamily membership, transmembrane domain prediction, and sequence length are included in an excel file available in the online Supplementary Material.

## ACKNOWLEDGEMENTS

This work was supported by National Science Foundation [Graduate Research Fellowship Program grant number DGE-1650441 to A.W.L., CAREER grant number DEB-1750394 to M.J.S.], National Institutes of Health [grant numbers DP2-GM128202 to M.J.S., S10OD018516 to the Cornell University Biotechnology Resource Center], and New York State Stem Cell Science [grant number CO29155 to Cornell BRC]. We thank Qi Sun and the Computational Biology Service Unit of the Cornell Life Sciences Core Laboratories Center for making software available on BioHPC and for providing helpful advice regarding gene annotation. We also thank Brook Luers and the Cornell Statistical Consulting Unit for statistics consultation.

## AUTHOR CONTRIBUTIONS

AWL and MJS conceptualized the study. AWL sequenced mRNA, manually annotated ORs, conducted all analyses, and wrote the paper. CMJ dissected and imaged antennal lobes, and wrote the Supplementary Materials & Methods section on Antennal Lobe Imaging. AWL and MFF labeled antennal lobe glomeruli. SEM and MJS sequenced and assembled the genomes. All authors helped edit the final manuscript.

