## Supplemental Materials & Methods, Tables S1-S3 and Figures S1-S9 for "Expansion and accelerated evolution of 9-exon odorant receptors in *Polistes* paper wasps"

| **CONTENTS** | **PAGE** |
|---|---|
| SUPPLEMENTARY MATERIALS & METHODS | 2 |
| • Antennal Lobe Imaging | 2 |
| • Antennal RNAseq | 3 |
| • Gene Annotation | 3 |
| • Phylogenetic Reconstruction | 6 |
| • Genomic Organization | 7 |
| • Sequence Analyses | 8 |
| REFERENCES | 9 |
| SUPPLEMENTARY TABLES | 10 |
| • Table S1: Summary Statistics for *Polistes* Genome Assemblies and Annotations | 10 |
| • Table S2: Summary Statistics of Antennal Transcriptome Assembly | 10 |
| • Table S3: Odorant Receptor Subfamily Size and Orthology Across *Polistes* | 11 |
| SUPPLEMENTARY FIGURES | 12 |
| • Figure S1: 3D Reconstruction of *P. fuscatus* Antennal Lobe | 12 |
| • Figure S2: Ideograms of 28 *P. fuscatus* genome scaffolds containing OR genes | 13 |
| • Figure S3: Maximum likelihood OR tree of five *Polistes* species | 14 |
| • Figure S4: Maximum likelihood OR tree of 9-exon ORs in five *Polistes* species | 16 |
| • Figure S5: Maximum likelihood OR tree of L subfamily ORs in five *Polistes* species | 18 |
| • Figure S6: Maximum likelihood OR tree of E subfamily ORs in five *Polistes* species | 20 |
| • Figure S7: Maximum likelihood OR tree of H subfamily ORs in five *Polistes* species | 22 |
| • Figure S8: Mean amino acid identity among orthologs in 9-exon vs. other subfamilies | 24 |
| • Figure S9: Portion of 9-exon tandem array *s13* with mapped antennal RNAseq reads | 25 |

### SUPPLEMENTARY MATERIALS & METHODS

#### Antennal Lobe Imaging

Immunocytochemistry: In the fall of 2018, 20 adult *P. fuscatus* wasps (10 male and 10 gyne) were collected in Ithaca, NY. Wasps were immobilized by cooling at 4°C, heads were removed, and brains were dissected from the head capsule and placed in ~1 ml of 4% paraformaldehyde in 0.01M phosphate buffered saline (PFA) overnight at 4°C. Mouse monoclonal anti-synapsin antibodies (SYNORF1, 3C11; Data Bank Hybridoma) were used as primary antibodies to label synaptically rich antennal lobe glomeruli. Phalloidin conjugated with TRITC (Tetramethylrhodamine Isothiocyanate; Invitrogen) was used to distinguish glomeruli borders and label antennal nerve tracts. F(ab')2 fragments of donkey anti-mouse antibodies conjugated to Alexa 488 were used as secondary antibodies to visualize anti-synapsin (Jackson ImmunoResearch Laboratories). After overnight fixation, brains were washed in 0.01M phosphate buffered saline with Triton-X 100 (PBS-TX) 6 times for 20 min. Brains were incubated for three nights at room temperature with anti-synapsin at 1:800 in PBS-TX solution. Brains were then washed again in PBS-TX (6 times for 20 min) and incubated for three nights at RT with donkey anti-mouse antibodies conjugated with Alexa 488 at 1:270, and then with Phalloidin-TRITC at 1:160 in PBS-TX. After staining with secondary antibodies, brains were post-fixed with 4% PFA for 10 min and dehydrated using increasing concentrations of methanol. Following full dehydration and two washes in 100% methanol, brains were cleared in methyl salicylate overnight. They were then mounted on slides in methyl salicylate.

Imaging: Images were collected using an inverted 880 confocal laser scanning microscope (Zeiss LSM880 Indimo, AxioObserver) with a C-Apochromat10x/0.45 water-immersion objective and a 1.8x digital zoom using the appropriate filter and laser setting for each fluorescent molecule and 8x line averaging. Image stacks were collected using 10-15 μm optical sections for both males and gynes. To individually label the glomeruli from confocal image stacks we used AVIZO and Amira software (Thermo Scientific, FEI) to make 3D reconstructions of the antennal lobe from whole mount preparations. Confocal stack files were imported into AVIZO software, and the voxel dimension outputs of each image stack were imported to provide correct dimensions (Thermo Scientific, FEI). Using the image segmentation function, we then identified glomeruli by hand throughout the entire image stack.

**Antennal RNAseq**

*P. fuscatus* mRNA library preparation and sequencing: In the fall of 2017, 12 adult *P. fuscatus* wasps (6 males and 6 gynes) from Ithaca, NY and Watkins Glen, NY were freeze-killed on dry ice and stored at -80°C. Each pair of antennae was cut at the base of the scape and weighed (paired gyne antennae = 1.27 mg ± 0.26 SD; paired male antennae = 1.57 mg ± 0.44 SD). RNA was isolated from 2 pooled antennae of each individual using the PureLink RNA Micro Kit (Ambion). Libraries were prepared using the NEBNext Ultra II RNA Library Prep Kit for Illumina (NEB #E7770L) with the NEBNext Poly(A) mRNA Magnetic Isolation Module (NEB #E7490) and NEBNext Multiplex Oligos for Illumina (NEB #E7600S). Libraries were analyzed on a fragment analyzer for quality control and sequenced on two lanes of an Illumina HiSeq (Illumina, San Diego, CA, USA).

Mapping mRNA reads of *P. fuscatus* and *P. dominula*: Raw reads from *P. fuscatus* antennae were trimmed using Trimmomatic v.0.39 to (1) remove adapter sequences, (2) cut leading and trailing bases below quality of 3, (3) cut 4-base wide windows when the average quality per base dropped below 20, and (4) remove reads below 35 base pairs (bp) long (Bolger et al. 2014). After trimming and filtering, more than 154 million paired end reads remained (75,846,826 male; 78,453,620 gyne) with an average length of 132 bp. Processed paired end reads were mapped, first to the *P. fuscatus* genome and subsequently to the genomes of two other *Polistes* species in the fuscopolistes clade (*P. metricus* and *P. dorsalis*), using STAR v2.6 to generate BAM files sorted by genome coordinate (Dobin et al. 2013). Each read was allowed to map to no more than one locus. Mapped reads from each sex were combined and assembled using Trinity genome-guided assembly (Haas et al. 2013). Summary statistics for the assembly are shown in Table S2. Raw reads from the whole-body transcriptome of a single individual *P. dominula* were downloaded from the Sequence Read Archive (Lopez-Osorio et al. 2017). To aid in manual annotation of ORs, reads were aligned to the *P. dominula* genome with STAR v2.7, using the same parameters as described above for *P. fuscatus*.

**Gene Annotation**

OR annotation in *P. fuscatus*: The *Polistes fuscatus* genome was assembled as described in Miller SE et al. 2020. Genome scaffolds containing putative ORs were identified based on sequence similarity with published insect ORs from a range of species: *Atta cephalotes*, *Acromyrmex echinatior*, *Apis mellifera*, *Camponotus floridanus*, *Cardiocondyla obscurior*, *Ceratosolen solmsi*,

*Drosophila melanogaster*, *Eulaema bombiformis*, *Euglossa dilemma*, *Euglossa flammea*, *Euglossa imperialis*, *Eulaema meriana*, *Eufriesea mexicana*, *Lasioglossum albipes*, *Microplitis demolitor*, *Monomorium pharaonis*, *Melipona quadrifasciata*, *Nasonia vitripennis*, *Solenopsis invicta* (Robertson et al. 2003, Robertson et al. 2010, Zhou et al. 2012, Zhou et al. 2015, Brand & Ramirez 2017, McKenzie & Kronauer 2018). The *P. fuscatus* genome was queried with amino acid sequences from each of the listed species using the TBLASTN algorithm with an e-value cutoff of 1e-5 (Altschul et al. 1997). Putative coding regions were identified by examining TBLASTN match regions and chaining the match regions according to criteria similar to those used by Zhou et al. (2012): neighboring match regions were included in the same coding region if the match regions were in close proximity and if the upstream amino acid query sequence was N-terminal to that of the downstream match region. Coding sequence (CDS) was determined by manually curating exon-intron boundaries in Geneious v11.1.5 using evidence from MAKER automated predictions, TBLASTN homology, Trinity predicted transcripts (aligned to genome using BLAT) and mapped RNAseq reads. We consulted version 24 of the Havana annotation guidelines to inform manual annotation of exon-intron splice sites (March 2016 guidelines by the Human and Vertebrate Analysis and Annotation group at WTSI). "AG" was the only permitted acceptor site. "GT" and "GC" were both acceptable donor sites, although "GC" was only present in a small number of genes. Amino acid sequences of ORs generated by this pipeline were used as queries in an iterative round of TBLASTN with scoring matrix PAM30 (Pearson 2013), to search for species-specific expansions, especially in the highly divergent 9-exon receptor clade.

OR annotation in 4 other *Polistes*: The *Polistes dorsalis* and *Polistes metricus* genomes were assembled and annotated as described in Miller SE et al. 2020. The *Polistes canadensis* and *Polistes dominula* genomes and annotations were accessed through NCBI (Patalano et al. 2015; Standage et al. 2016). Putative ORs were identified by using TBLASTN with the same sample of Hymenopteran protein query sequences used during manual annotation of *P. fuscatus* ORs, with the addition of *P. fuscatus* OR protein sequences. Uncertain gene models were aligned using Muscle version 3.8.425 with maximum 4 iterations and gene models were manually adjusted based on homology with functional ORs in *P. fuscatus* (Edgar 2004). Gene models were called pseudogenes if they exhibited frame-shift mutation, premature stop codons, or unacceptable 5' donor or 3' receptor splice sites.

Naming ORs: All ORs were named with a species-specific prefix using the first letter of the genus followed by the first three letters of the species epithet (e.g. PfusOR#). Orco was not numbered (e.g. PfusOrco), and no OR was labeled OR1, since some previous studies refer to Orco as OR1. All tuning ORs were numbered in the order in which they appeared in the genome, beginning with OR2. Therefore, OR names with the same number do not necessarily indicate orthology among *Polistes*. Incomplete gene models were named with a suffix to indicate which features of the gene model are missing, following Oxley et al. (2014): "NTE" missing N-terminus, "INT" missing interior sequence, "CTE" missing C terminus, "NI" missing N terminus and interior sequence, "NC" missing N and C termini. All pseudogenes were given the suffix "PSE."

Search for ORs in unassembled genomic reads: Genome misassembly is a potential cause of overestimation of pseudogenes, and is more common in repetitive regions of the genome. For example, many ORs that were previously annotated as pseudogenes in the clonal raider ant were not identified in a new, chromosome arm level assembly of the genome, which also yielded many more functional OR gene models (McKenzie & Kronauer 2018). Given the completeness of the *Polistes* genomes (Table S1) and the close correspondence between functional *P. fuscatus* ORs and number of *P. fuscatus* antennal lobe glomeruli, this type of error likely accounts for minimal annotation errors in this species. We searched for sequence similarity to ORs in the unassembled genomic reads of *P. dorsalis*, the species in which we initially found the fewest functional ORs. The 10x Genomics Supernova Assembler version 2.1.1 failed to assemble unmapped *P. dorsalis* genomic reads greater than or equal to 150 nucleotides (~21.4 million paired-end reads). Therefore, the reads themselves were formatted as a blast database and queried using TBLASTN at an e-value cutoff of 1e-5. The same sample of Hymenopteran protein sequences used during manual annotation, with the addition of the putatively functional OR protein sequences of all five *Polistes* species, were used to search the unaligned *P. dorsalis* genomic reads. This resulted in significant hits to 120 paired-end reads. Given that on average 159 ± 555 paired-end reads map to a given OR gene in *P. fuscatus*, it is unlikely that many ORs were lost in unassembled genomic reads. However, it cannot be ruled out that genome misassembly may account for overestimation of pseudogenes in some cases.

Quality assessment of putatively functional ORs: All *Polistes* OR protein sequences greater than 300 amino acids in length were analyzed using InterProScan version 5.17-56.0 to check for the InterPro olfactory receptor annotation IPR004117, the 7tm odorant receptor Pfam accession

PF02949, and the odorant receptor PANTHER accession PTHR21137 (Jones P et al. 2014). For each of 192 orthologous groups predicted by OrthoFinder, the orthologs were aligned using Muscle version 3.8.425 with maximum 4 iterations, and alignments were inspected for anomalies that could be the result of annotation error. Adjustments were made to the gene models in 29 orthologous groups. The majority of these adjustments were changes in the length of already annotated exons with few or no mapped mRNA reads.

**Phylogenetic Reconstruction**

Hymenoptera-wide analysis of OR gene family: We used published OR sequences from *Apis mellifera*, *Nasonia vitripennis* (Robertson & Wanner 2006; Robertson et al. 2010), and *Camponotus floridanus* (Zhou et al. 2012), along with OR sequences of *P. fuscatus* (this study) to reconstruct the phylogeny of the OR gene family. All amino acid (aa) sequences of chemoreceptors at least 300 aa in length were aligned using MAFFT version 7.453 with the high accuracy "E-INS-i" method ("genafpair" option combined with 1000 cycles of iterative refinement ("maxiterate")) (Katoh and Standley 2013). The alignment was trimmed using trimAl v3 with the "automated1" option to automatically choose the parameters that best fit the qualities of the alignment (Capella-Gutierrez et al. 2009), generating a phylip file with 1066 sequences of 231 aa length. Gene trees were generated with RAxML version 8.2.12 using the Maximum Likelihood optimality criterion with 100 rapid bootstrap inferences, CAT model of rate heterogeneity across sites (Stamatakis 2006), JTT amino acid replacement matrix, F option to use empirical base frequencies from the input alignment ("-m PROTCATJTTF"), "-p 12345" to set initial random number for parsimony search, *N. vitripennis* Orco as outgroup, "-f a" to combine bootstrap and maximum likelihood searches in one run, "-x 12345" to set initial random number for bootstrap search, "-N 100" to perform a bootstrap analysis informed by 100 unique maximum likelihood trees, and "-k" to add branch lengths to trees (Stamatakis 2014, Jones DT et al. 1992). The tree was rerooted in FigTree v1.4.4 to visually clarify the position of the Orco gene clade (available online at https://github.com/rambaut/figtree/releases/tag/v1.4.4). The tree was visualized in R version 3.6.1 using ggtree (Figure 4A; Yu et al. 2017; R Core Team 2019).

Estimation of gene gain and loss events: Gene duplication and loss events were reconstructed by reconciling a gene tree with a species tree in NOTUNG version 2.9.1.3 (Chen et al. 2000; Durand et al. 2006; Vernot et al. 2008). A gene tree of putatively functional OR proteins at least 300 amino acids in length was generated using published protein sequences for 4 ants (*A.*

*echinatior*, *A. cephalotes*, *S. invicta*, *C. floridanus*; Zhou et al. 2012, 2015), 2 bees (*A. mellifera*, *L. albipes*; Robertson et al. 2010, Zhou et al. 2015), 3 nonsocial wasps (*N. vitripennis*, *M. demolitor*, *C. solmsi*; Robertson et al. 2010, Zhou et al. 2015), and 5 social wasps (*P. fuscatus*, *P. metricus*, *P. dorsalis*, *P. canadensis*, *P. dominula*; annotated in this study). The gene tree was generated as described above for the Hymenoptera OR tree, without rerooting. Following Zhou et al. 2012, all internal branches with bootstrap support values below 70 were collapsed into polytomies in R using ggtree. Notung was run in rearrangement mode using an edge weight (bootstrap value) threshold of 90, with cost of "1.5" for a duplication and cost of "1" for a loss.

Analysis of *Polistes* OR gene family: The OR sequences of *P. fuscatus*, *P. metricus*, *P. dorsalis*, *P. canadensis*, and *P. dominula* were filtered to include only ORs of at least 300 amino acids in length. These were aligned using MAFFT ("E-INS-i" method) and trimmed in trimal using the parameters described above for the Hymenoptera OR tree. The tree was generated using RAxML, rerooted in FigTree, and visualized in R using ggtree as described above. Trees in the supplementary materials were visualized using Interactive Tree Of Life (iTOL) version 5.6.2 (Letunic & Bork 2019). Orthology of *Polistes* OR amino acid sequences was inferred using sequence similarity and gene collinearity in the genome. Orthologs were called based on similarity of amino acid sequences with OrthoFinder version 2.2.7 using DIAMOND for sequence similarity searches with an e-value cutoff of 0.001 (Emms & Kelly 2015). OrthoFinder predicted 135 orthologous groups that included gene copies from both *P. fuscatus* and *P. dorsalis*. For the purpose of pairwise $d_N/d_S$ analysis, this set of orthologs was expanded by inspecting the gene tree (Figure 5A; Figure S3) and Muscle alignments (see "Genomic organization" below) of syntenic regions. Gene copies within a clade supported by a bootstrap value of at least 70 and showing synteny were considered orthologs in the expanded ortholog set of 150 single copy orthologs between *P. fuscatus* and *P. dorsalis*.

*Polistes* species phylogeny: The phylogenetic tree of *Polistes* species in Figure 1 was built from Supplemental Fig 2 in Sheehan et al. 2015 using the "physketch" function in the phytools R package (Revell 2012). The initial tree was constructed using the 16S ribosomal RNA gene and the cytochrome oxidase subunit I gene (Sheehan et al. 2015).

**Genomic Organization**

Two functional ORs separated by a pseudogenized OR were considered neighbors if their coding sequences were within 5kb of each other. Neighboring OR protein sequences were aligned using

Muscle v3.8.425 with maximum 4 iterations to determine percent amino acid identity. One-way ANOVA comparing percent amino acid identity of neighbors across eight tandem arrays was performed using the "aov" function, and Tukey HSD test with the "TukeyHSD" function, both of which are in the base R package stats version 3.6.1 (R Core Team 2019).

**Sequence Analyses**

Tests for positive selection: All putatively functional *Polistes* ORs greater than 350 amino acids in length were used in analyses of episodic diversifying positive selection within OR subfamilies using aBSREL in HyPhy version 2.5.15 (Smith MD et al. 2015; Pond et al. 2020). Subfamily-specific alignments were generated using MAFFT version 7.453 ("E-INS-i") and trimmed in trimal with the "gappyout" option. Trees were generated using RAxML as above, except the "PROTGAMMAAUTO" model was used to automatically choose the best protein model given likelihood. CDS alignments were generated using trimal with the "backtranslate" option added to the above parameters. The significance threshold for each aBSREL analysis was Holm-Bonferroni corrected p=0.05. Values of pairwise $d_N/d_S$ for orthologs shared by *P. fuscatus* and *P. dorsalis* were calculated using PAML version 4.9 (Yang 2007) program yn00 with the Yang & Nielsen (2000) method. Values of $d_N$, $d_S$, and omega were compared between 9-exon and non-9-exon orthologous pairs using the function "t.test" in the base R package stats version 3.6.1.

Test for gene conversion: To search for gene conversion events, all *Polistes* OR subfamily codon alignments were analyzed using GENECONV version 1.81a with options "/lp" to list all pairwise significant fragments, "/w123" to set initial random number, and "-nolog" to avoid a segmentation fault in UNIX (Sawyer 1989). Bonferroni-corrected KA p-values below 0.05 were considered significant.

### SUPPLEMENTARY TABLES

**Table S1**: Summary Statistics for *Polistes* Genome Assemblies and Annotations

| Species | Size[a] | N50[b] | Genes[c] | BUSCO euk.[d] | BUSCO arth.[e] | CEGMA euk.[f] | Reference[g] |
|---|---|---|---|---|---|---|---|
| *P. fuscatus* | 219 | 9.11 | 12034 | 97.7% (296/303) | 99.4% (1060/1066) | N/A | Miller et al. 2020 (GCA_010416935.1) |
| *P. metricus* | 220 | 4.63 | 12032 | 98.0% (297/303) | 100% (1066/1066) | N/A | Miller et al. 2020 (GCA_010416925.1) |
| *P. dorsalis* | 209 | 5.37 | 11311 | 98.0% (297/303) | 99.8% (1064/1066) | N/A | Miller et al. 2020 (GCA_010416905.1) |
| *P. canadensis* | 211 | 0.52 | 16997 | N/A | N/A | 98.8% (245/248) | Patalano et al. 2015 (GCA_001313835.1) |
| *P. dominula* | 208 | 1.63 | 12153 | N/A | N/A | 99.2% (246/248) | Standage et al. 2016 (GCA_001465965.1) |

a. Size in Mb of genome assembly
b. Scaffold N50 in Mb
c. Total number of annotated genes in genome
d. Annotation completeness as measured by BUSCO score for core eukaryotic genes
e. Annotation completeness as measured by BUSCO score for core arthropod genes
f. Annotation completeness as measured by CEGMA score for core eukaryotic genes
g. Reference to genome publication followed by the GenBank assembly accession

**Table S2**: Summary Statistics of Antennal Transcriptome Assembly

| | | | |
|---|---|---|---|
| Total trinity 'genes' | 163537 | | |
| Total trinity transcripts | 211491 | | |
| Percent GC | 33.41 | Based on longest isoform per 'gene' | |
| contig N10 | 13311 | contig N10 | 7460 |
| contig N20 | 8798 | contig N20 | 4062 |
| contig N30 | 6356 | contig N30 | 2259 |
| contig N40 | 4547 | contig N40 | 1383 |
| contig N50 | 3026 | contig N50 | 953 |
| Median contig length | 463 | Median contig length | 399 |
| Average contig | 1152.76 | Average contig | 703.89 |
| Total assembled bases | 243799125 | Total assembled bases | 115111742 |

**Table S3**: Odorant Receptor Subfamily Size and Orthology Across *Polistes*

| Subfam. | *P. fus.* | *P. met.* | *P. dor.* | *P. can.* | *P. dom.* | proportion 1:1 orthology[a] | ortholog identity[b] |
|---|---|---|---|---|---|---|---|
| 9-exon | 105 | 109 | 85 | 101 | 92 | 32 | 91 ± 4.8 |
| A | 1 | 1 | 1 | 1 | 1 | 100 | 85.6 |
| B | 1 | 1 | 1 | 1 | 1 | 100 | 95.9 |
| C | 1 | 1 | 1 | 1 | 1 | 100 | 96.6 |
| D | 1 | 1 | 1 | 0 | 1 | 0 | NA |
| E | 8 | 7 | 6 | 6 | 5 | 63 | 94.5 ± 2.7 |
| F | 5 | 5 | 5 | 4 | 5 | 80 | 96.6 ± 1.5 |
| G | 2 | 2 | 2 | 1 | 2 | 0 | NA |
| H | 9 | 8 | 5 | 6 | 6 | 33 | 93.3 ± 0.5 |
| I | 1 | 1 | 1 | 1 | 1 | 100 | 95.2 |
| J | 4 | 4 | 4 | 4 | 4 | 100 | 96.2 ± 1.4 |
| K | 1 | 1 | 1 | 1 | 1 | 100 | 98.4 |
| L | 25 | 25 | 25 | 23 | 22 | 60 | 94.9 ± 2.7 |
| M | 1 | 1 | 1 | 1 | 1 | 100 | 92.5 |
| O | 3 | 3 | 3 | 3 | 4 | 67 | 92.1 ± 2.4 |
| P | 1 | 1 | 1 | 1 | 1 | 100 | 94.9 |
| Q | 5 | 7 | 8 | 7 | 6 | 80 | 94.7 ± 1.5 |
| R | 1 | 1 | 1 | 1 | 1 | 100 | 95.3 |
| T | 14 | 14 | 14 | 14 | 14 | 71 | 94.91 ± 3.1 |
| V | 6 | 6 | 6 | 6 | 6 | 100 | 96.2 ± 1.2 |
| W | 1 | 1 | 1 | 1 | 1 | 100 | 92 |
| X | 1 | 1 | 1 | 1 | 1 | 100 | 97.3 |
| Z | 1 | 1 | 1 | 1 | 1 | 100 | 95.5 |
| Orco | 1 | 1 | 1 | 1 | 1 | 100 | 99.9 |
| Unclass. | 1 | 1 | 1 | 1 | 1 | 100 | 92.3 |
| All | 200 | 204 | 177 | 188 | 180 | 50 | 93.6 ± 4 |

a. proportion of *P. fuscatus* ORs with 1:1 orthology across all species
b. mean pairwise amino acid identity among OrthoFinder 1:1 orthologs (%) ± SD

### SUPPLEMENTARY FIGURES

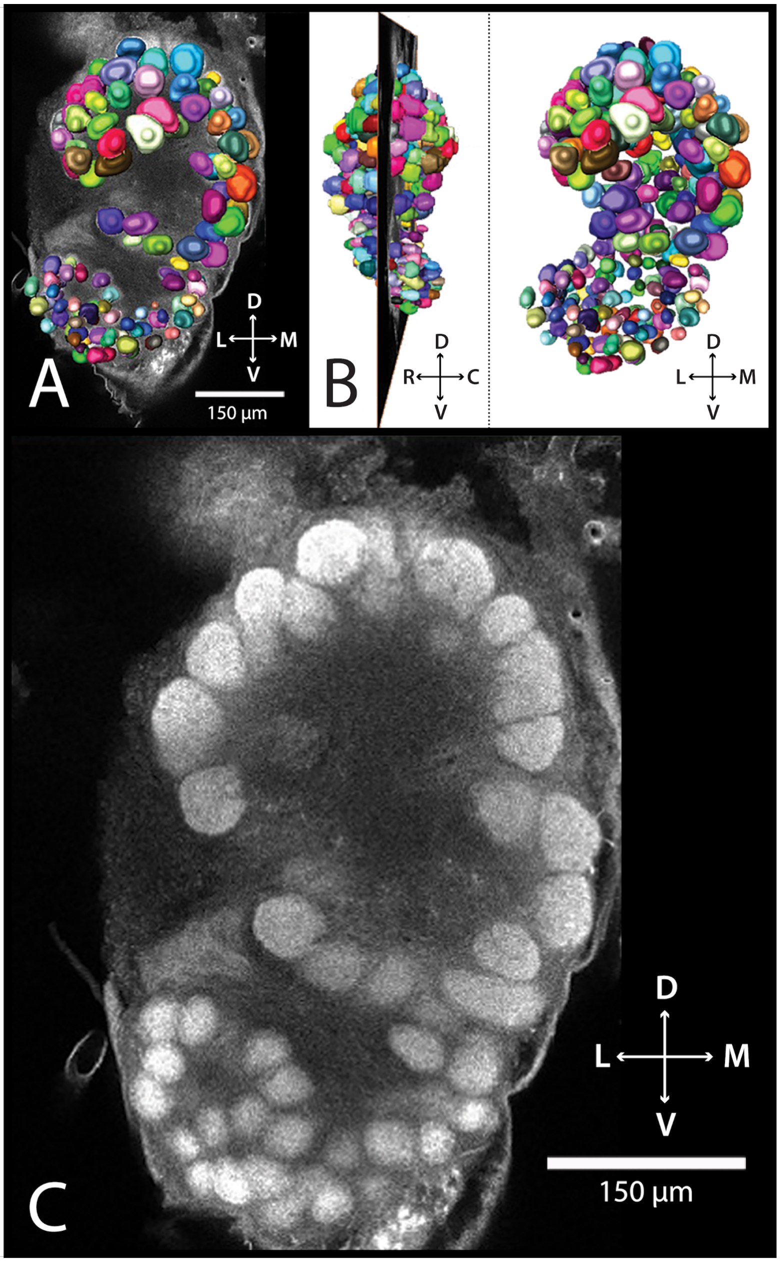


**Figure S1**: (A) 3D reconstruction of the anti-synapsin stained left antennal lobe of a *P. fuscatus* gyne superimposed on an orthogonal slice of a confocal image stack. (B) Sagittal view of 3D reconstruction with orthogonal slice corresponding to (A), accompanied by frontal view on the right. Colors denote individual glomeruli. (C) Confocal microscopy slice. Scale bars represent 150 µm. D: dorsal; V: ventral; M: medial; L: lateral; R: rostral; C: caudal.

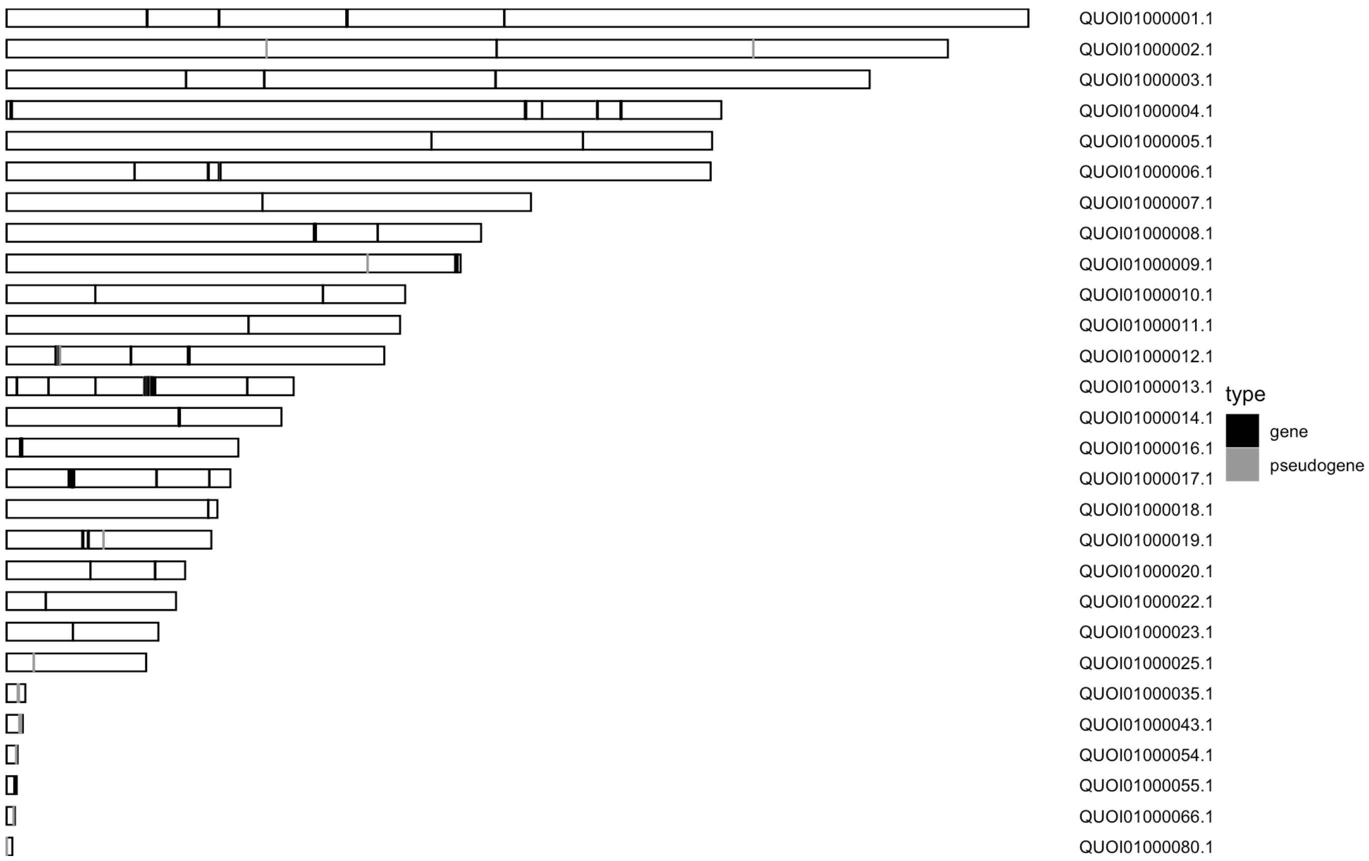


**Figure S2**: Ideograms of 28 *Polistes fuscatus* reference genome scaffolds which contain OR genes (black) and OR pseudogenes (gray) represented by vertical lines. The last unique digits of each scaffold name correspond to the scaffold numbers (*s#*) referred to in the main text (e.g. "QUOI01000017.1" corresponds to the main text scaffold "*s17*").

Figure S3: Maximum likelihood OR tree of five Polistes species

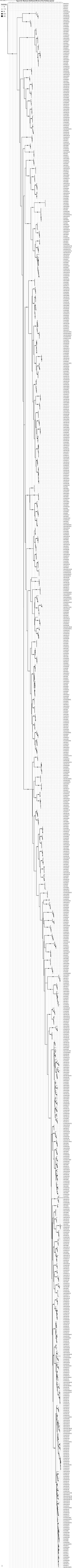

**Figure S3**: (see preceding page) Maximum likelihood OR protein tree of five *Polistes* species (Pfus: *P. fuscatus*; Pmet: *P. metricus*; Pdor: *P. dorsalis*; Pcan: *P. canadensis*; Pdom: *P. dominula*). Branches with bootstrap support values greater than or equal to 70 are labeled with a black circle. Scale bar represents mean amino acid substitutions per site.

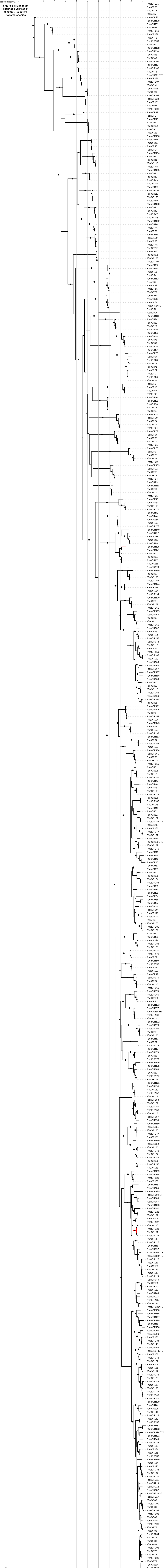


**Figure S4: Maximum likelihood OR tree of 9-exon ORs in five Polistes species**

**Figure S4**: (see preceding page) Maximum likelihood OR protein tree of 9-exon subfamily ORs in five *Polistes* species. Three branches that were found by HyPhy aBSREL to display dN/dS > 1 indicating positive selection are highlighted in red. Branches with bootstrap support values greater than or equal to 70 are labeled with a black circle. Scale bar represents mean amino acid substitutions per site.

**Figure S5: Maximum likelihood OR tree of L subfamily ORs in five Polistes species**

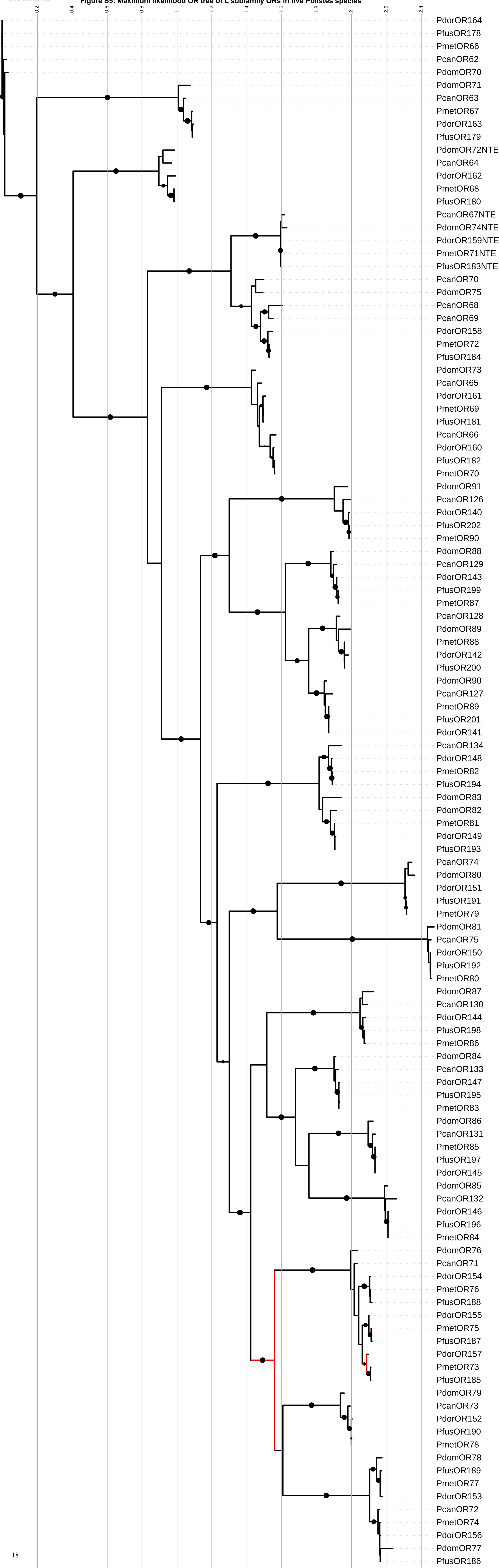

**Figure S5**: (see preceding page) Maximum likelihood OR protein tree of L subfamily ORs in five *Polistes* species. Three branches that were found by HyPhy aBSREL to display dN/dS > 1 indicating positive selection are highlighted in red. Branches with bootstrap support values greater than or equal to 70 are labeled with a black circle. Scale bar represents mean amino acid substitutions per site.

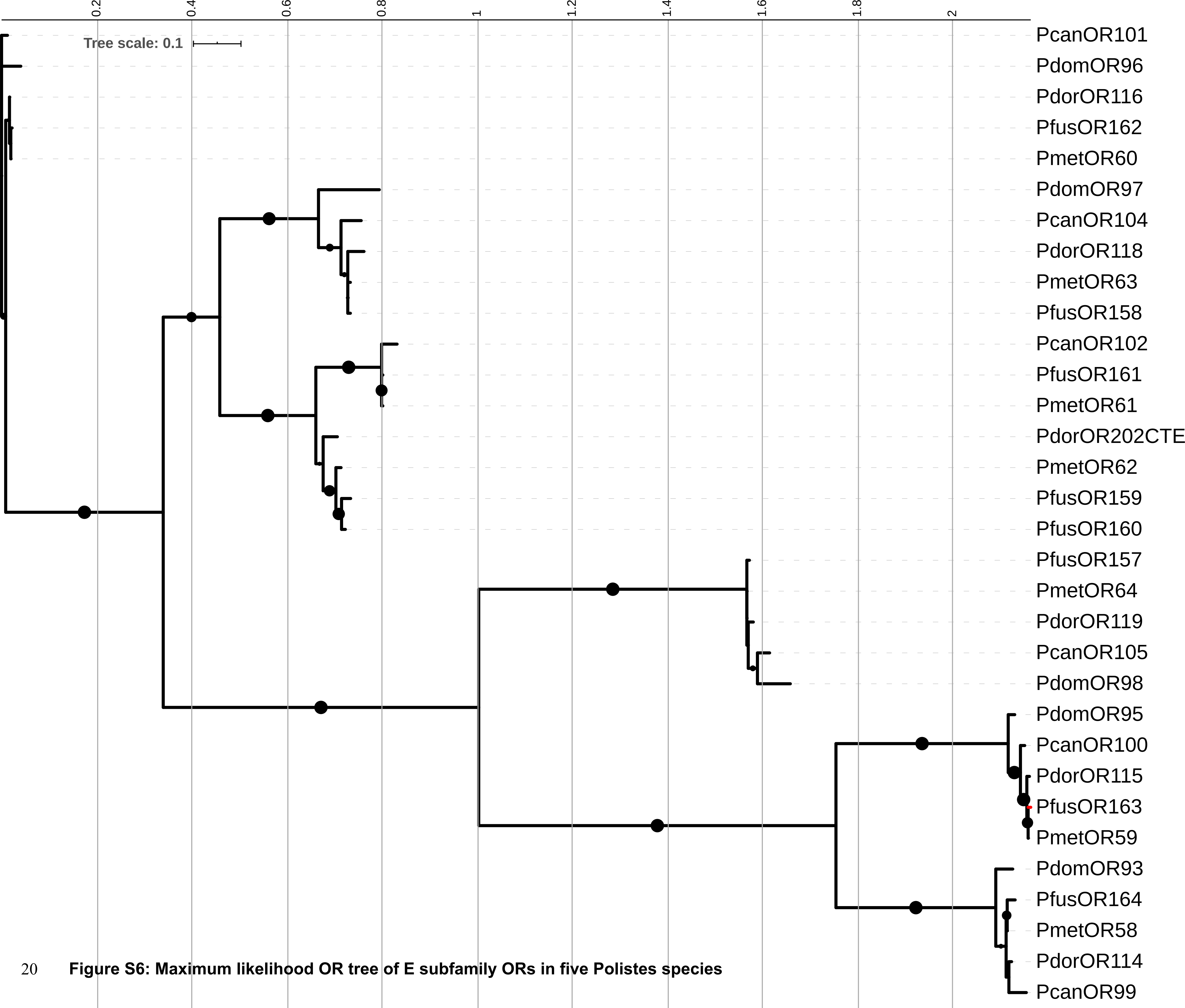


**Figure S6: Maximum likelihood OR tree of E subfamily ORs in five Polistes species**

**Figure S6**: (see preceding page) Maximum likelihood OR protein tree of E subfamily ORs in five *Polistes* species. Three branches that were found by HyPhy aBSREL to display dN/dS > 1 indicating positive selection are highlighted in red. Branches with bootstrap support values greater than or equal to 70 are labeled with a black circle. Scale bar represents mean amino acid substitutions per site.

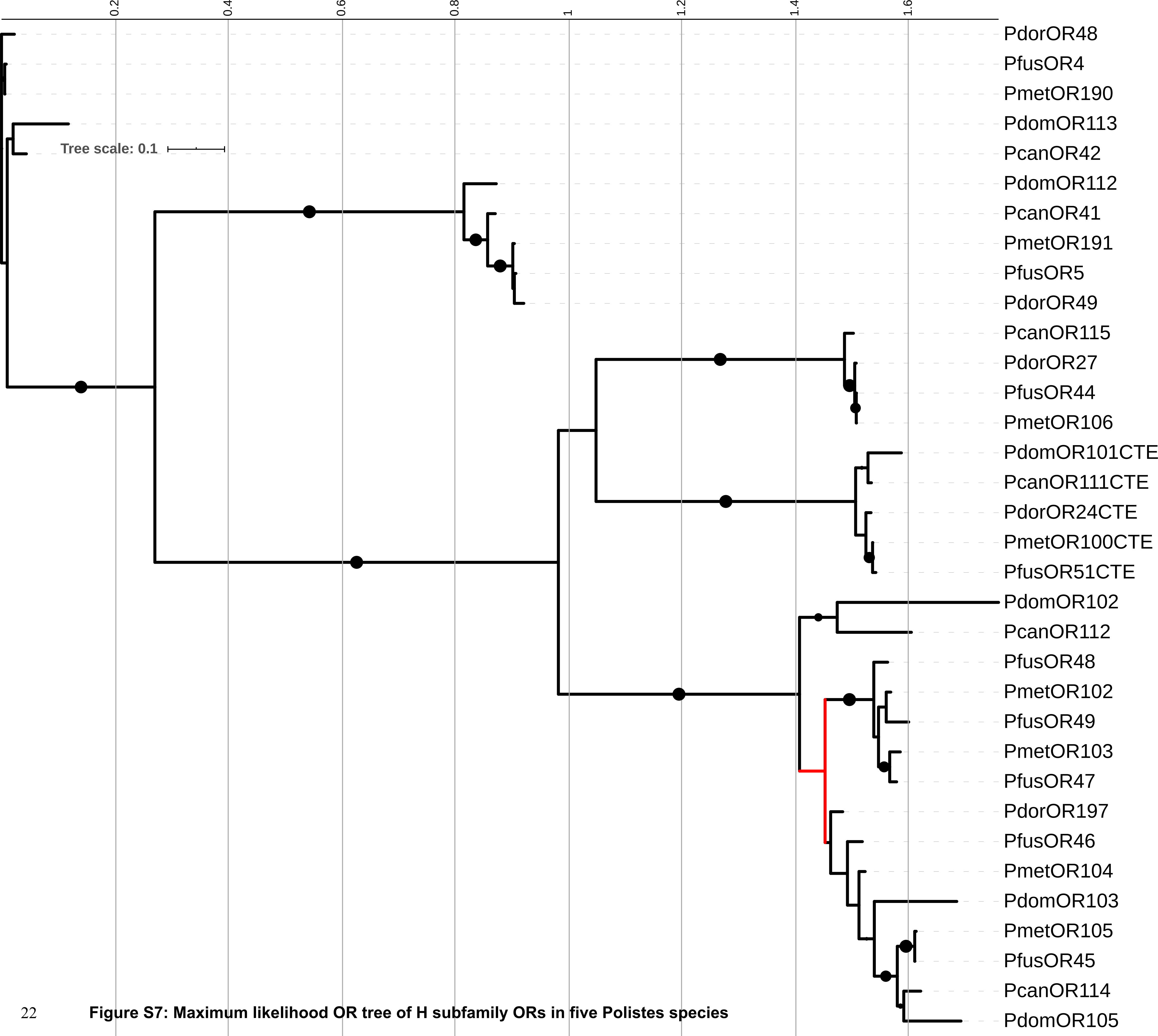


**Figure S7: Maximum likelihood OR tree of H subfamily ORs in five Polistes species**

**Figure S7**: (see preceding page) Maximum likelihood OR protein tree of H subfamily ORs in five *Polistes* species. Three branches that were found by HyPhy aBSREL to display dN/dS > 1 indicating positive selection are highlighted in red. Branches with bootstrap support values greater than or equal to 70 are labeled with a black circle. Scale bar represents mean amino acid substitutions per site.

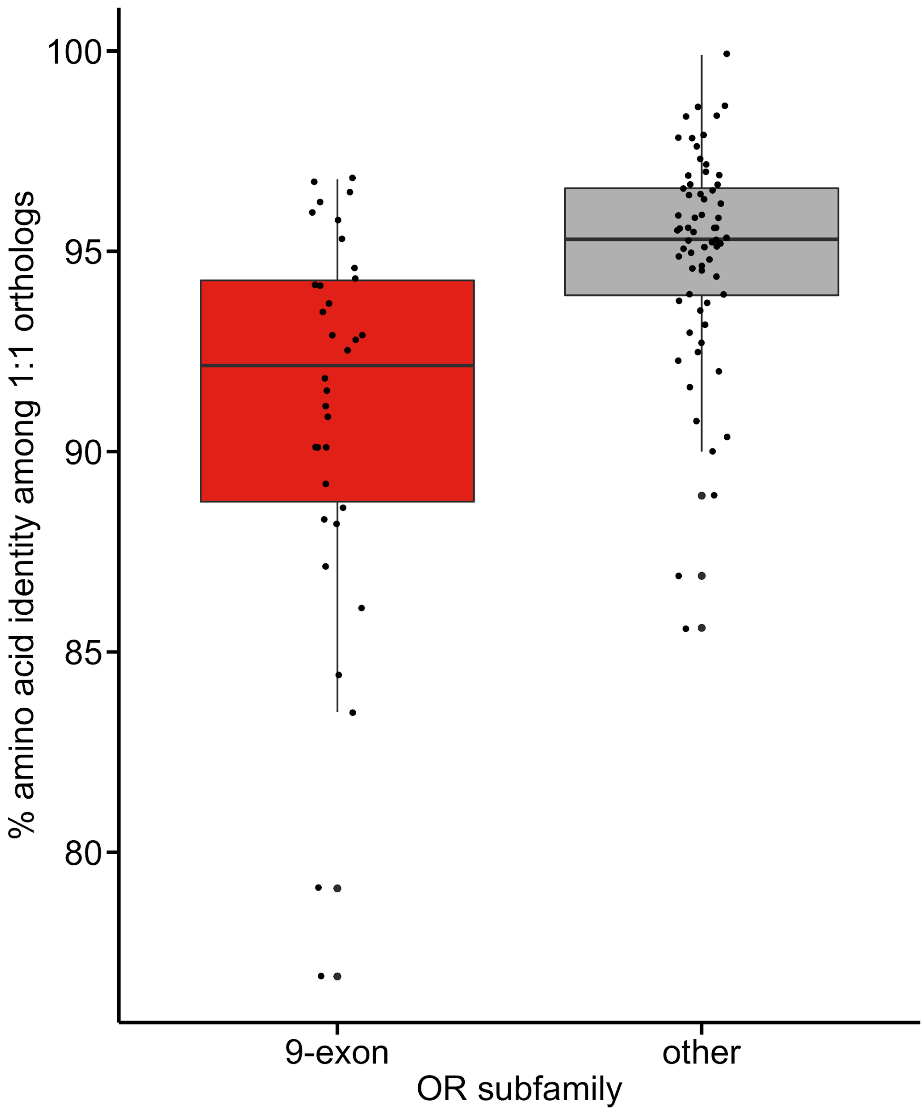


**Figure S8**: The 91% mean amino acid identity among 1:1 9-exon subfamily orthologs in *Polistes* (red) is significantly lower than the 95% mean amino acid identity among 1:1 orthologs in all other OR subfamilies (gray) (Welch Two Sample t-test: P = 7.051e-05).

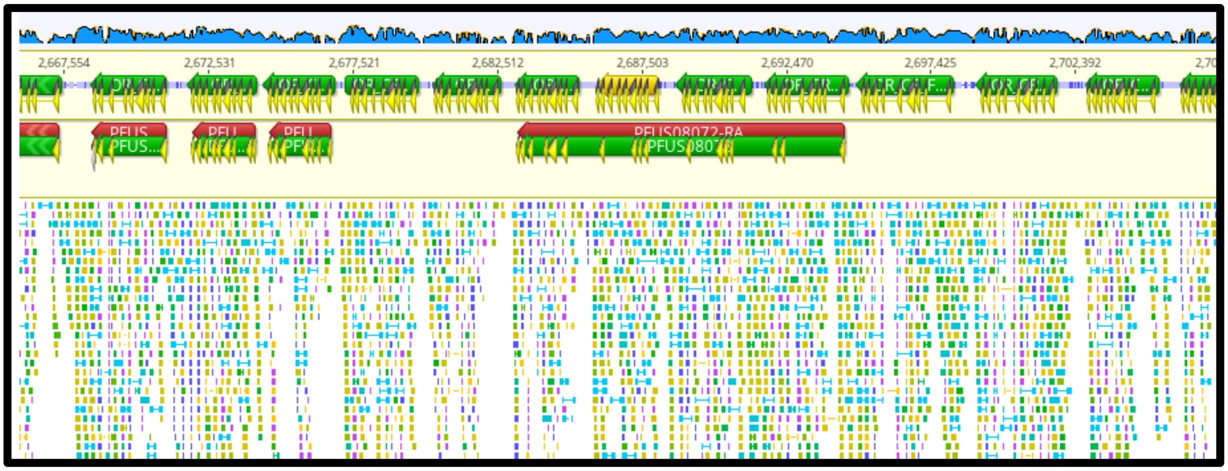


**Figure S9**: Screenshot of Geneious genome browser view of a portion of 9-exon tandem array *s13* highlighting the usefulness of antennal RNAseq reads in OR gene annotation. The blue coverage map represents the RNAseq reads, aligned on the bottom track. The top annotation track shows manually annotated OR genes (green = putatively functional; orange = pseudogenes) and the bottom annotation track shows the automated annotations generated by the MAKER pipeline, which was otherwise highly effective in annotating most genes (BUSCO 97.7% eukaryotic genes, Table S1).
